# YAP-controlled MHC-I and CXCL10 expression potentiates antitumor immunity independently of IFNγ signaling

**DOI:** 10.1101/2023.02.25.530003

**Authors:** Linyuan Peng, Liang Zhou, Huan Li, Desheng Lu, Minxian Qian, Zhongyuan Wang

## Abstract

Many efforts are underway to improve immune checkpoint blockade (ICB) therapy including potentiating MHC-I antigen processing and presentation. Accumulating evidence links the Hippo pathway to immunotherapy response, but the understanding of how the tumor-intrinsic Hippo signaling regulating antitumor immunity is limited. Here, we report that inhibition of the Hippo pathway coactivator YAP in tumor cells increases the expression of genes involving in MHC-I antigen processing and presenting machinery (APM) and CXCL10, promoting robust tumor-infiltrating of cytotoxic CD8^+^ T lymphocytes (CTLs) and overcoming tumor resistance to anti-PD1 ICB therapy. Mechanically, we find that YAP/TEAD complex directly binds and recruits NuRD complex to the NLRC5 promoter to repress NLRC5 transcription, thereby blunting MHC-I antigen presentation. Patient cohort analysis revealed that YAP expression negatively correlated with the expression of NLRC5 and MHC-I APM and intratumoral infiltration of CTLs. Collectively, our results suggest that a novel tumor-promoting function of YAP depends on NLRC5 to impair MHC-I antigen processing and presentation and provide a rationale for inhibiting YAP activity in ICB therapy for cancer.

## Introduction

Immune checkpoint blockade (ICB) has revolutionized cancer treatment in clinical studies (*1–4*). However, the response varies depending on tumor types, and only a subset of patients benefits from ICB (*5*). Comprehensive profiling of the tumor-intrinsic mechanisms of heterogeneous responses should provide opportunities to improve ICB efficacy. Multiple mechanisms have been identified, including the degree of intra-tumoral infiltration and activation of immune cells, especially cytotoxic CD8^+^ T lymphocytes (CTLs) which correlates positively with the success of ICB (*6–9*). CTLs specifically recognized tumor cells through directly binding of the T cell receptor (TCR) to neo-antigens in an MHC class I (MHC-I) antigen processing and presenting machinery (APM)-dependent manner (*10*).

Antigen processing and presenting process involves the generation and loading of peptides on MHC-I molecules, which are composed of a transmembrane heavy chain association with β2 microglobulin (B2M) and expressed on the surface of all nucleated cells. The process initiates with proteins degradation by proteasome, especially immunoproteasome to generate MHC-I peptides. Once transported into the endoplasmic reticulum (ER) by transporter associated with antigen processing (TAP), the peptides are loaded on nascent MHC-I molecules with the aid of peptide loading complex (PLC). Then the peptide loaded MHC-I molecules are exported to the cell membrane and presented to CTLs (*11, 12*). The reduced expression of MHC-I APM genes by genomic mutations or epigenetic and/or transcriptional silencing dampens antigen presentation, TCL recognition and activation and antitumor activity (*13*). Loss of MHC-I expression is frequently observed in several tumor types and associated with unfavorable disease outcomes (*14–18*). NLRC5, also known as MHC-I transactivator (CITA), is an important gene expression regulator of MHC-I APM genes encoding MHC-I heavy or light chains (MHC-I and B2M), immunoproteasome components (PSMB8 and PSMB9) and peptide transporter associated with antigen processing (TAP1) (*19*). Reduced expression or activity of NLRC5 due to epigenetic alterations, copy number loss, or somatic mutations results in decreased MHC-I APM gene expression, CTL activation and patient prognosis (*20*). However, the mechanisms controlling the expression of MHC-I APM genes in cancer are still elusive.

The Hippo pathway, first identified in Drosophila melanogaster, plays critical regulatory functions in disease development, particularly tumorigenesis in mammals. The canonical mammalian Hippo pathway consists of a tumor suppressor gene Neurofibromatosis type 2 (NF2, also known as merlin, an essential component of the Hippo cascade), core kinases MST1/2 and LATS1/2, and downstream effectors YAP and TAZ (*21*). The MST1/2 phosphorylate and activate Lats1/2, which in turn negatively regulate YAP and TAZ by phosphorylation. Shuttle between the cytoplasm and the nucleus, YAP and TAZ function as transcriptional co- activators to bind to transcriptional factors such as TEAD1-4 to regulate gene expression, which promotes organ size during development, and tumorigenesis in adult tissues (*22*). More recently, the role of the Hippo pathway in regulating antitumor immunity has received increasing attention. The Hippo pathway regulates the development and functionality of T cells, B cells and macrophages, which is crucial for tumor immunity. Furthermore, the deregulation of tumor-intrinsic Hippo pathway enables tumor cells to avoid immune surveillance (*23, 24*). However, the crosstalk between the Hippo pathway in tumor cells and the immune microenvironment remains to be fully investigated.

Here we investigated a tumor-intrinsic role of YAP in regulating the expression of MHC-I APM genes. YAP/TEAD complex binds NLRC5 promoter at specific sites and blocks its transcription, and consequently decreases MHC-I antigen processing and presenting, thereby contributing to tumor immune evasion and play crucial roles in response to ICB.

## Results

YAP or TEAD loss augments NLRC5 and MHC-I APM gene expression To explore the role of Hippo pathway in antitumor immunity, we knocked down Yap in two malignant mouse cancer cell lines (4T1 and MC38) using lentiviral mediated shRNA (Fig. 1, C and D, and fig. S2, C and D). Yap knockdown (KD) cells were inoculated into severe immunodeficient NCG (NOD/ShiLtJGpt-*Prkdc*^em26Cd52^*Il2rg*^em26Cd22^/Gpt) mice or syngeneic immunocompetent mice. Knockdown of Yap induced a reduction in tumor growth by about 40% in NCG mice, however, the tumor growth of Yap KD cells in syngeneic immunocompetent mice was attenuated by more than 60% (fig. S1, A and C). Accordingly, knockdown of Yap led to a much more obvious survival benefit in syngeneic immunocompetent mice than in NCG mice (fig. S1, B and D). These observations suggested that YAP has a tumor-intrinsic effect in tumor development via regulation of immune microenvironment.

**Fig. 1.**
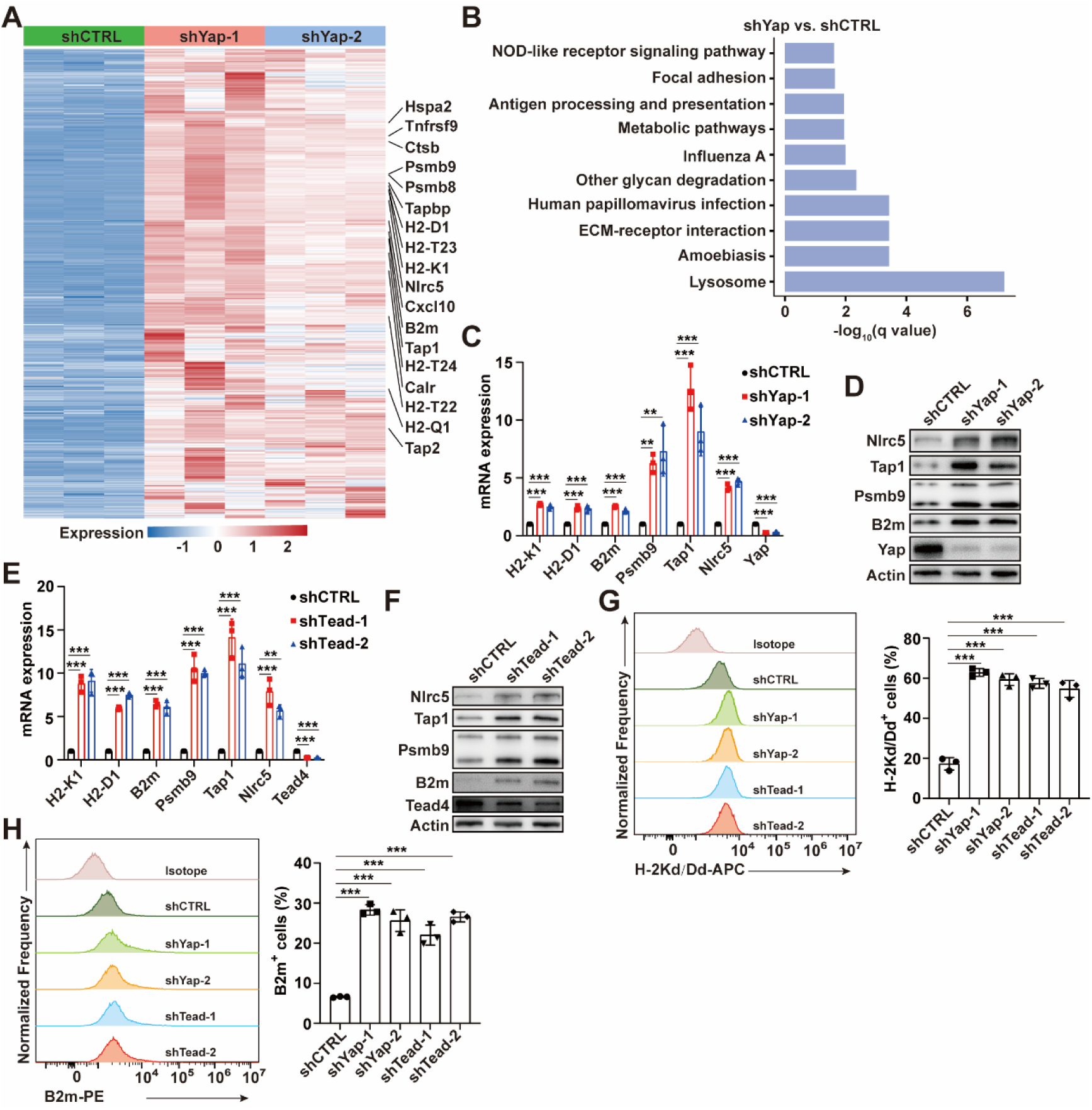
YAP or TEAD deficiency upregulates the expression of NLRC5 and MHC-I APM genes. **(A)** Heat map of differential expression of genes induced by Yap knockdown (KD) in 4T1 cells by RNA-seq. **(B)** KEGG analyses show the altered pathways after Yap depletion. **(C)** qPCR analysis of the mRNA expression of MHC-I APM genes and Nlrc5 gene in 4T1 cells with or without Yap KD (n = 3). **(D)** Immunoblot (IB) analysis of 4T1 cells with or without Yap KD. **(E)** qPCR analysis of the mRNA expression of MHC-I APM genes and Nlrc5 gene in 4T1 cells with or without Tead2/3/4 KD (n = 3). **(F)** IB analysis of 4T1 cells with or without Tead2/3/4 KD. **(G)** Flow cytometry detection of cell-surface level of H- 2Kd/Dd in 4T1 cells with or without Yap or Tead2/3/4 KD. The percentage of H-2Kd/Dd positive cells is shown (n = 3). **(H)** Flow cytometry detection of cell-surface level of B2m in 4T1 cells with or without Yap or Tead2/3/4 KD. The percentage of B2m positive cells is shown (n = 3). Data are presented as means ± SD. *P* values were calculated by one-way ANOVA in C and E. *P* values in G and H were determined by unpaired two-sided *t*- test. \*\**P* < 0.01, \*\*\**P* < 0.001.

To elucidate the molecular basis by which YAP regulated immune microenvironment, we performed RNA-seq on wild-type Yap (Yap-WT) or Yap-KD 4T1 cells. RNA-seq results revealed that multiple MHC-I APM components including those encoding MHC-I heavy or light chains (H2- K1, H2-D1 and B2m), immunoproteasome components (Psmb8 and Psmb9) and peptide transporter associated with antigen processing (Tap1) were upregulated in Yap-KD cells (Fig. 1A). Interestingly, MHC-I transactivator Nlrc5 which regulates the transcription of MHC-I APM genes and the proinflammatory chemokines Cxcl10 were also upregulated (Fig. 1A). The Kyoto Encyclopedia of Genes and Genomes (KEGG) analysis showed that antigen processing and presentation genes were enriched in YAP-KD cells (Fig. 1B). Likewise, gene set enrichment analysis (GSEA) revealed that the expression of MHC-I APM genes were enhanced after YAP KD (fig. S2, A and B).

To validate RNA-seq results, qPCR analysis was performed in three mouse cancer cell lines (4T1 and MC38). We found either Yap or Tead KD increased the expression of MHC-I APM genes encoding MHC-I transactivator Nlrc5, H2-K1, H2-D1, B2m, PSMB8, PSMB9 and TAP1 (Fig. 1, C and E, and fig. S2, C and E). Consistently, western blotting showed the protein expression of Nlrc5, B2m, Psmb9 and Tap1 were upregulated in either Yap or Tead KD cells (Fig. 1, D and F, and fig. S2, D and F); flow cytometry revealed the cell-surface expression of the MHC-I heavy chains (H-2Kd/Dd and H-2Kb for 4T1 and MC38 cells respectively) were higher after Yap or Tead KD (Fig. 1G and fig. S2G), with similar expression observed for cell-surface B2m in 4T1 and MC38 cells (Fig. 1H and fig. S2H). Next, the analysis was extended in two human cancer cell lines (MDA-MB-231 and SUM159) and observed both YAP and TEAD negatively regulated the mRNA and protein expression of MHC-I transactivator NLRC5, MHC-I heavy or light chains (HLA-A, HLA-B, HLA-C and B2M), PSMB9 and TAP1 as well as cell-surface levels of HLA and B2M (fig. S3, A to L). Consistent with our data, the mRNA level of YAP correlated negatively with the expression of NLRC5, HLA-A, B2M, PSMB9 and TAP1 in five human malignancies (fig. S3M).

### The Hippo pathway is involved in regulation of NLRC5 and MHC-I APM gene expression

To further investigate whether the upstream Hippo pathway components modulating the expression of NLRC5 and MHC-I APM genes, tumor suppressor gene NF2, core kinases MST2 or LATS1 was transfected into NF2 deficient MDA-MB-231 cells (*25*). Overexpression of wild-type NF2, MST2 and LATS1, but not patient-derived inactive mutant NF2 (L64P) upregulated the mRNA and protein expression of NLRC5, HLA-A/B/C, B2M, PSMB9 and TAP1 as well as cell-surface levels of HLA and B2M (Fig. 2, A to D, G and H). Next, we found a TEAD-binding deficient YAP (S94A) protein increased while an active mutant YAP (5SA) protein reduced the mRNA and protein expression of NLRC5, HLA-A/B/C, B2M, PSMB9 and TAP1 as well as cell-surface levels of HLA and B2M (Fig. 2, E to H, and fig. S4, A to D). It is recognized that ECM stiffness regulates YAP activity, we monitored if the expression of NLRC5 and MHC-I APM genes in cells grown on fibronectin-coated acrylamide hydrogels of varying stiffness from 1 to 40 kPa (*25*). As expected, growing cells on soft matrices (1 kPa) suppressed YAP activity indicated by YAP regulated gene CTGF (fig. S4E). Consistently, the mRNA and protein expression of NLRC5, HLA-A/B/C, B2M, PSMB9 and TAP1 as well as cell-surface levels of HLA and B2M were induced by growing cells on soft matrices (1 kPa). Moreover, a selective inhibitor of YAP/TEAD transcriptional activity CA3 promoted the mRNA and protein expression of Nlrc5, H2-K1, H2-D1, B2M, PSMB9 and TAP1 as well as cell-surface levels of H-2Kd/Dd and B2m (Fig. 2, I to L). Taken together, we conclude that the Hippo pathway inactivation or YAP activation impaired the expression of NLRC5 and MHC-I APM genes.

**Fig. 2.**
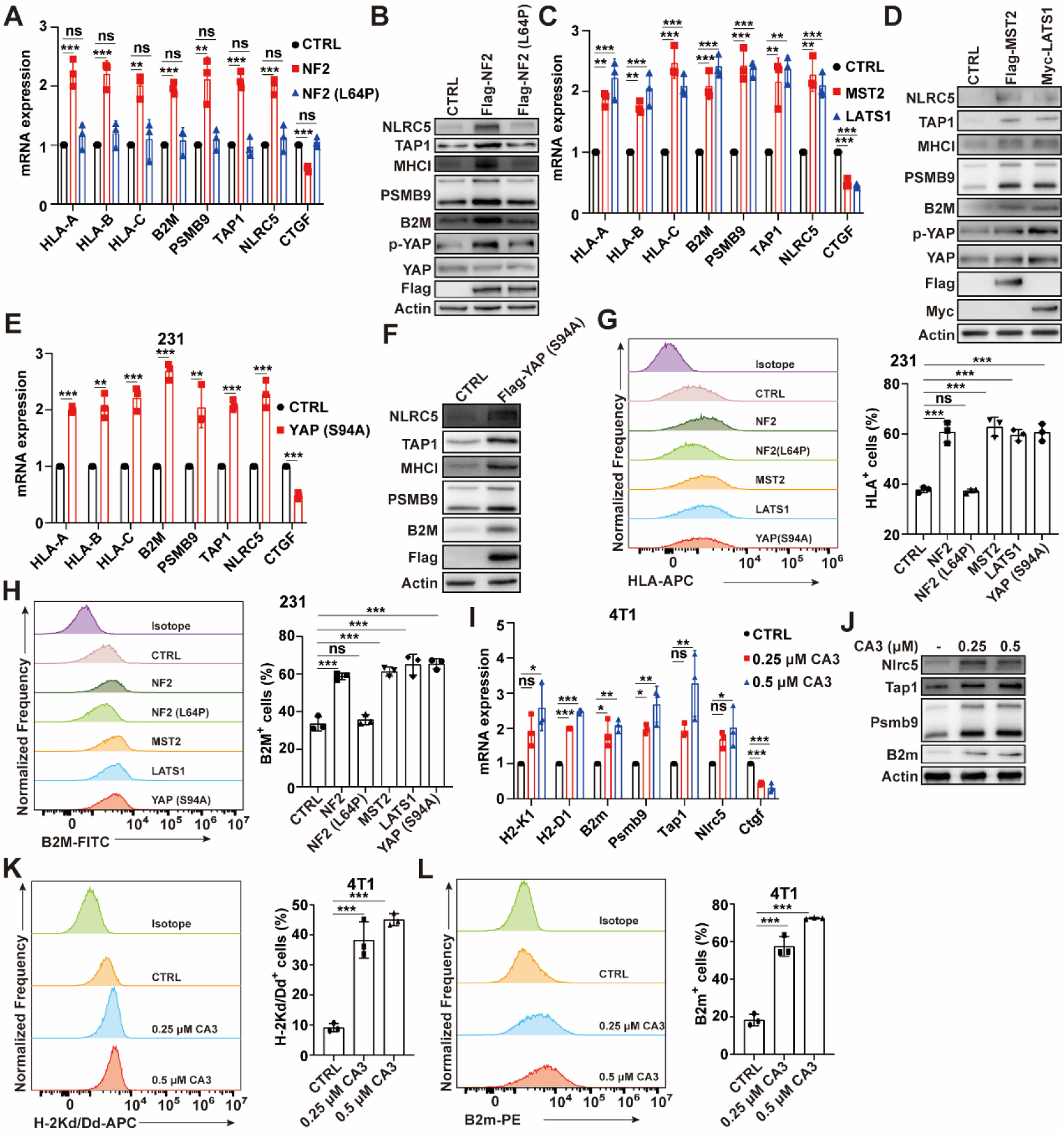
The Hippo pathway regulates the expression of NLRC5 and MHC-I APM genes. **(A)** qPCR analysis of the mRNA expression of MHC- I APM genes and NLRC5 gene in MDA-MB-231 cells transfected with wild type NF2 or missense patient mutation NF2 (L64P). A Hippo target gene CTGF is used as positive control (n = 3). **(B)** Immunoblot (IB) analysis of MDA-MB-231 cells transfected with wild type NF2 or missense patient mutation NF2 (L64P). **(C)** qPCR analysis of the mRNA expression of MHC-I APM genes and NLRC5 gene in MDA-MB-231 cells transfected with Flag-MST2 or Myc-LATS1 plasmid. CTGF is used as positive control (n = 3). **(D)** IB analysis of MDA-MB-231 cells transfected with Flag-MST2 or Myc-LATS1 expression plasmid. **(E)** qPCR analysis of the mRNA expression of MHC-I APM genes and NLRC5 gene in MDA- MB-231 cells transfected with or without a TEAD-binding deficient YAP (S94A) expression plasmid. CTGF is used as positive control (n = 3). **(F)** IB analysis of MDA-MB-231 cells transfected with or without a TEAD- binding deficient YAP (S94A) expression plasmid. **(G)** Flow cytometry detection of cell-surface level of HLA-A/B/C in MDA-MB-231 cells transfected with NF2, NF2 (L64P), MST2, LATS1 or YAP (S94A). The percentage of HLA-A/B/C positive cells is shown (n = 3). **(H)** Flow cytometry detection of cell-surface level of B2M in MDA-MB-231 cells transfected with NF2, NF2 (L64P), MST2, LATS1 or YAP (S94A). The percentage of B2M positive cells is shown (n = 3). **(I)** qPCR analysis of the mRNA expression of MHC-I APM genes and Nlrc5 gene in 4T1 cells treated with or without the indicated YAP/TEAD inhibitor CA3. Ctgf is used as positive control (n = 3). **(J)** IB analysis of 4T1 cells treated with or without the indicated CA3. **(K)** Flow cytometry detection of cell-surface level of H-2Kd/Dd in 4T1 cells treated with or without the indicated CA3. The percentage of H-2Kd/Dd positive cells is shown (n = 3). **(L)** Flow cytometry detection of cell-surface level of B2m in 4T1 cells treated with or without the indicated CA3. The percentage of B2m positive cells is shown (n = 3). Data are presented as means ± SD. *P* values were calculated by one-way ANOVA in A, C and I. *P* values in E, G, H, K and L were determined by unpaired two-sided *t*-test. \*\**P* < 0.01, \*\*\**P* < 0.001. ns, not significant.

### The YAP/TEAD complex directly binds and recruits NuRD complex to NLRC5 promoter to represses NLRC5 transcription

Considering NLRC5 as MHC-I transactivator and our observations that YAP or TEAD KD induced the expression of NLRC5 and MHC-I APM genes, we predicted the YAP/TEAD complex may regulate NLRC5 transcription and surveyed both human and mouse NLRC5 promoter regions by MatInspector (Genomatix). Intriguingly, human NLRC5 promoter harbors two putative TEAD binding sites (“CATTCC”, nominated as TB1 and TB2), while one putative TEAD binding site (TB) motif was located in mouse Nlrc5 promoter around the transcription start site (TSS) (fig. S5). Genomes of other species were also surveyed and found the NLRC5 TB motifs are conserved in primates, rodents and artiodactyla (fig. S5). Furthermore, we analyzed the TEAD4 chromatin immunoprecipitation coupled with high-throughput sequencing (ChIP–seq) dataset (GSM1010868) and found the potential TEAD4-binding peaks in the NLRC5 gene locus. To further validate the physical association of the YAP/TEAD complex with the NLRC5 promoter, ChIP-qPCR assays using anti-YAP or anti-TEAD4 antibody exhibited the enrichment of endogenous YAP and TEAD4 near the TB sequences of NLRC5 in both human and mouse cells (Fig. 3, A and B).

**Fig. 3.**
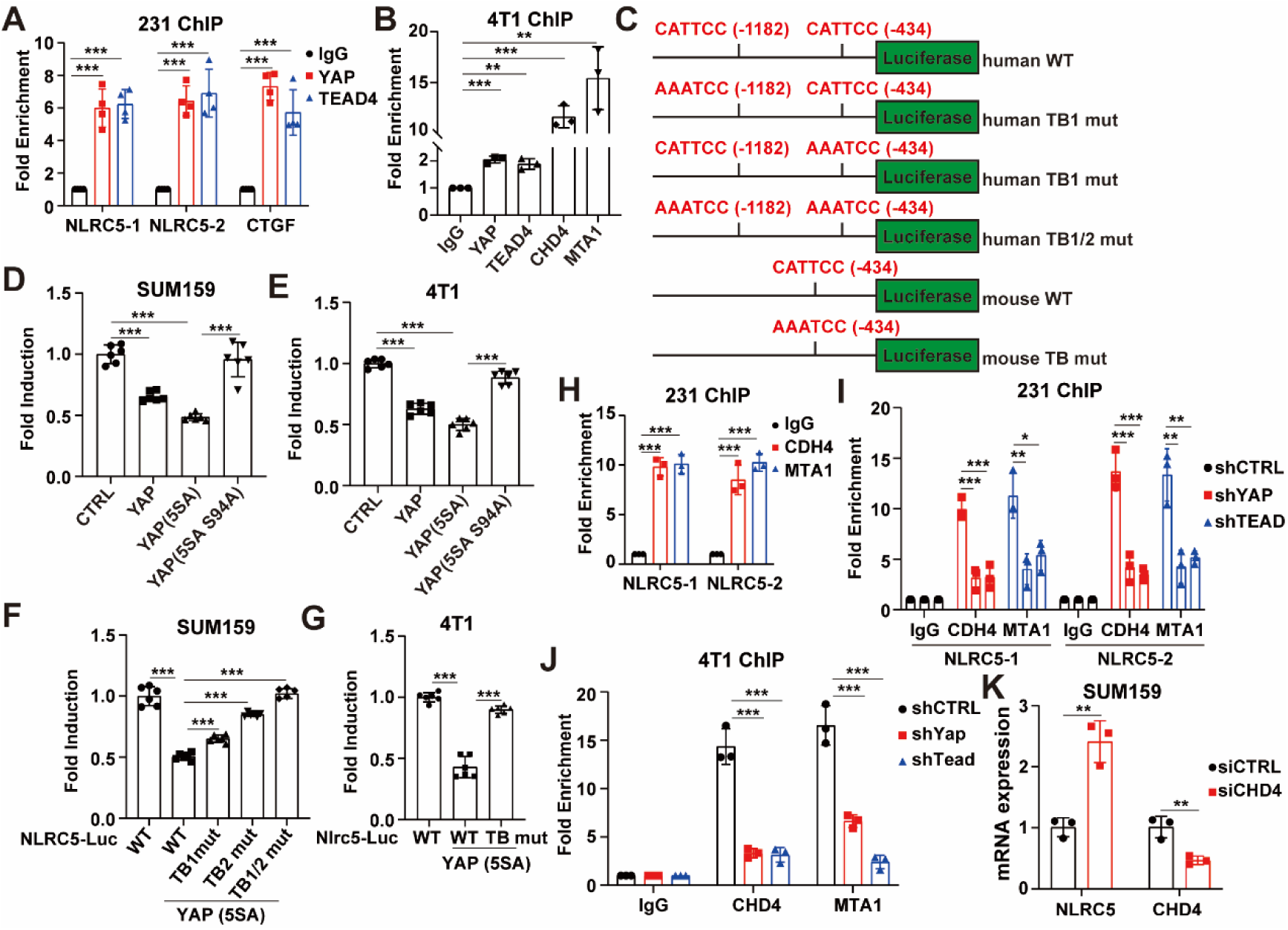
NLRC5 is the direct target of YAP/TEAD complex. **(A)** ChIP- qPCR analysis of YAP and TEAD4 signals using primers specific to NLRC5 promoter in MDA-MB-231 cells. YAP/TEAD target gene CTGF is used as positive control (n = 4). **(B)** ChIP-qPCR analysis of YAP, TEAD4, CHD4 and MTA1 signals using primers specific to Nlrc5 promoter in 4T1 cells (n = 3). **(C)** Schematic diagram showing the human and murine NLRC5 reporter constructs used in this study. The TEAD binding (TB) motifs and their positions with respect to the TSS are indicated in red. The numbers in parentheses refer to the positions of the first base of each TB motif. **(D)** Luciferase activity of human NLRC5 reporter in SUM159 cells transfected with the indicated expression plasmids (n = 6). **(E)** Luciferase activity of murine Nlrc5 reporter in 4T1 cells transfected with the indicated expression plasmids (n = 6). **(F)** Luciferase activity of human wild type or TB mutant NLRC5 reporter in SUM159 cells transfected with YAP (5SA) expression plasmids (n = 6). **(G)** Luciferase activity of murine wild type or TB mutant NLRC5 reporter in 4T1 cells transfected with YAP (5SA) expression plasmids (n = 6). **(H)** ChIP-qPCR analysis of CHD4 and MTA1 signals using primers specific to NLRC5 promoter in MDA-MB-231 cells (n = 3). **(I)** ChIP-qPCR analysis of CHD4 and MTA1 signals using primers specific to NLRC5 promoter in MDA-MB-231 cells with YAP or TEAD1/3/4 KD (n = 3). **(J)** ChIP-qPCR analysis of CHD4 and MTA1 signals using primers specific to Nlrc5 promoter in 4T1 cells with YAP or TEAD2/3/4 KD (n = 3). **(K)** qPCR analysis of the mRNA expression of NLRC5 and CHD4 in SUM159 cells transfected with or without siCHD4. Data are presented as means ± SD. *P* values were calculated by one-way ANOVA in A, H, I and J. *P* values in B, D, E, F, G and K were determined by unpaired two-sided *t*-test. \*\**P* < 0.01, \*\*\**P* < 0.001.

To further verify that NLRC5 is a directly repressed target gene of the YAP/TEAD complex, both human and mouse wild-type NLRC5 promoters (-2000 to -1 bp) and TB mutation promoters were separately cloned and inserted upstream of the luciferase gene (Fig. 3C). The NLRC5 reporter activity was efficiently reduced in wild-type YAP or constitutively active YAP (5SA), but not TEAD-binding deficient YAP (5SA S94A) overexpressing cells, confirming the requirement of TEAD (Fig. 3, D and E). When one of the two TB motifs was mutated in human NLRC5 promoter, the reporter activity was still repressed by YAP (5SA) (Fig. 3F). However, the disruption of both TB sequences in human NLRC5 promoter or the only TB sequence in mouse NLRC5 promoter largely abrogated this suppressive effect (Fig. 3, F and G). Together, the YAP/TEAD complex is involved in the repression of NLRC5 transcription.

The repressor activity of the YAP/TEAD complex has been reported to be mediated by the NuRD complex in which CHD4 is a chromatin remodeling subunit and MTA sububits (MTA1/2/3) mediate NuRD complex targeting to the YAP/TEAD complex (*26*). Indeed, using ChIP-qPCR, CHD4 and MTA1 enriched near the TB sequences of NLRC5 (Fig. 3, B and H), while the occupancy was decreased by YAP or TEAD depletion (Fig. 3, I and J).

Blocking the activity of NuRD by siCHD4 rescued NLRC5 expression in YAP (5SA) overexpressing cells (Fig. 3K). These results indicate that the NuRD complex mediates the repressor function of the YAP/TEAD complex.

### The YAP/TEAD complex abolishes the expression of MHC-I APM genes through NLRC5

Next, we investigated whether the YAP/TEAD complex-repressed MHC-I APM gene expression depends on NLRC5. Using qPCR, western blotting and flow cytometry, YAP or TEAD depletion enhanced the mRNA and protein expression of NLRC5 and MHC-I APM genes as shown previously, whereas simultaneously depletion of NLRC5 in YAP or TEAD KD cells reversed the increased effects (Fig. 4 and fig. S6).

**Fig. 4.**
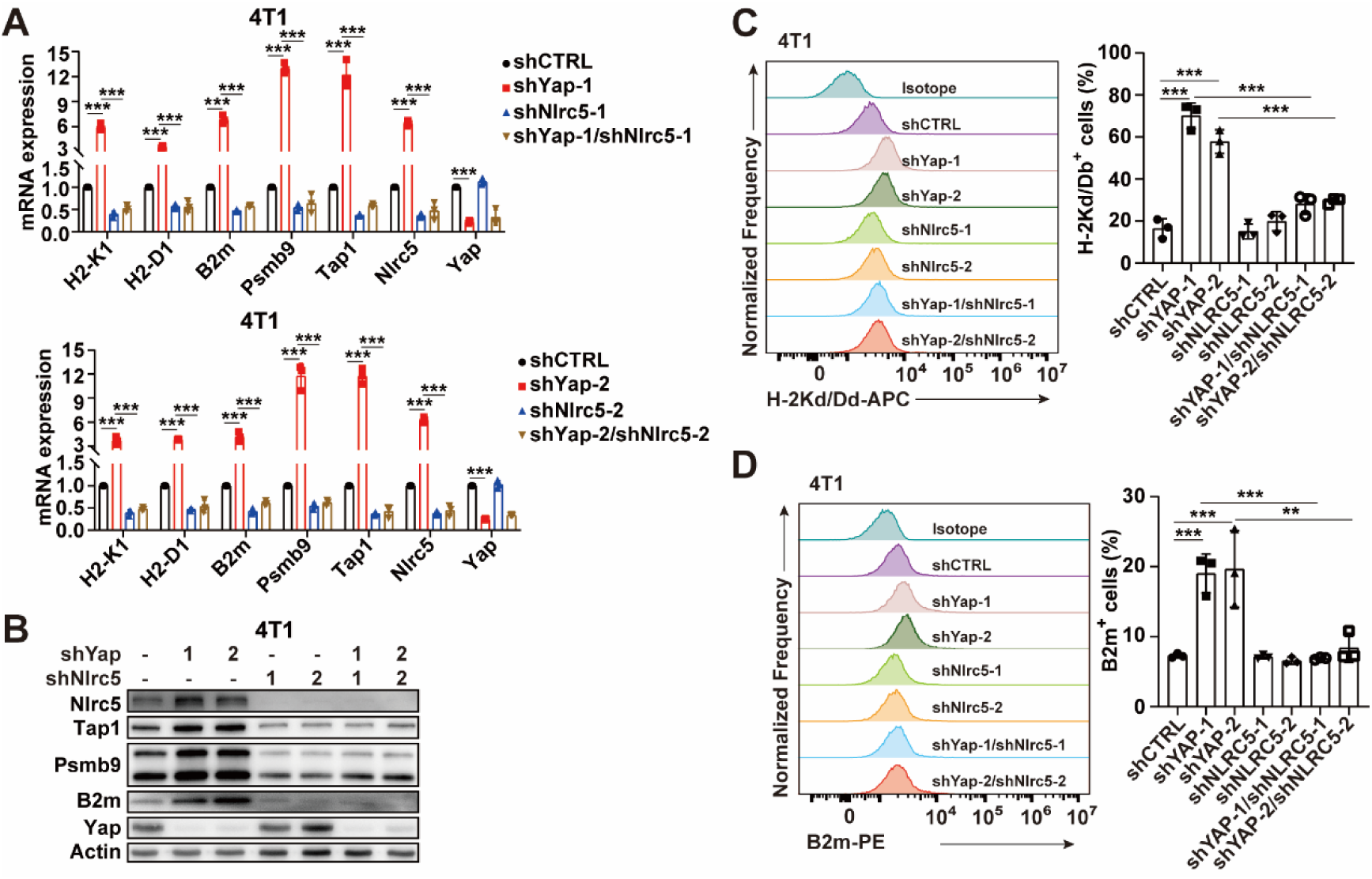
YAP deficiency promotes the expression of MHC-I APM genes through NLRC5. **(A)** qPCR analysis of the mRNA expression of MHC-I APM genes and Nlrc5 gene in 4T1 cells infected with two independent shYap and/or shNlrc5 lentivirus (n = 3). **(B)** Immunoblot (IB) analysis of 4T1 cells infected with two independent shYap and/or shNlrc5 lentivirus. **(C)** Flow cytometry detection of cell-surface level of H-2Kd/Dd in 4T1 cells infected with two independent shYap and/or shNlrc5 lentivirus. The percentage of H-2Kd/Dd positive cells is shown (n = 3). **(D)** Flow cytometry detection of cell-surface level of B2m in 4T1 cells infected with two independent shYap and/or shNlrc5 lentivirus. The percentage of B2m positive cells is shown (n = 3). Data are presented as means ± SD. *P* values were calculated by one-way ANOVA in A and B. *P* values in C and D were determined by unpaired two-sided *t*-test. \*\**P* < 0.01, \*\*\**P* < 0.001.

### The YAP/TEAD complex disrupts CXCL10 expression

As reported previously, Cxcl10 secreted by cancer cells attracts CTLs homing in the tumor microenvironment (*27*). Our RNA-seq data showed that the chemokine Cxcl10 was upregulated in Yap KD cells (Fig. 1A). To verify RNA-seq results, we performed qPCR analysis and found depletion of YAP or TEAD induced an increase in CXCL10 expression in both human (MDA-MB-231 and SUM159) and mouse (4T1 and MC38) cancer cell lines (Fig. 5, A and B, and fig. S7, A and B). Consistently, ELISA revealed that the protein expression of CXCL10 was efficiently promoted after YAP or TEAD KD (Fig. 5, C and D, and fig. S7, C and D).

**Fig. 5.**
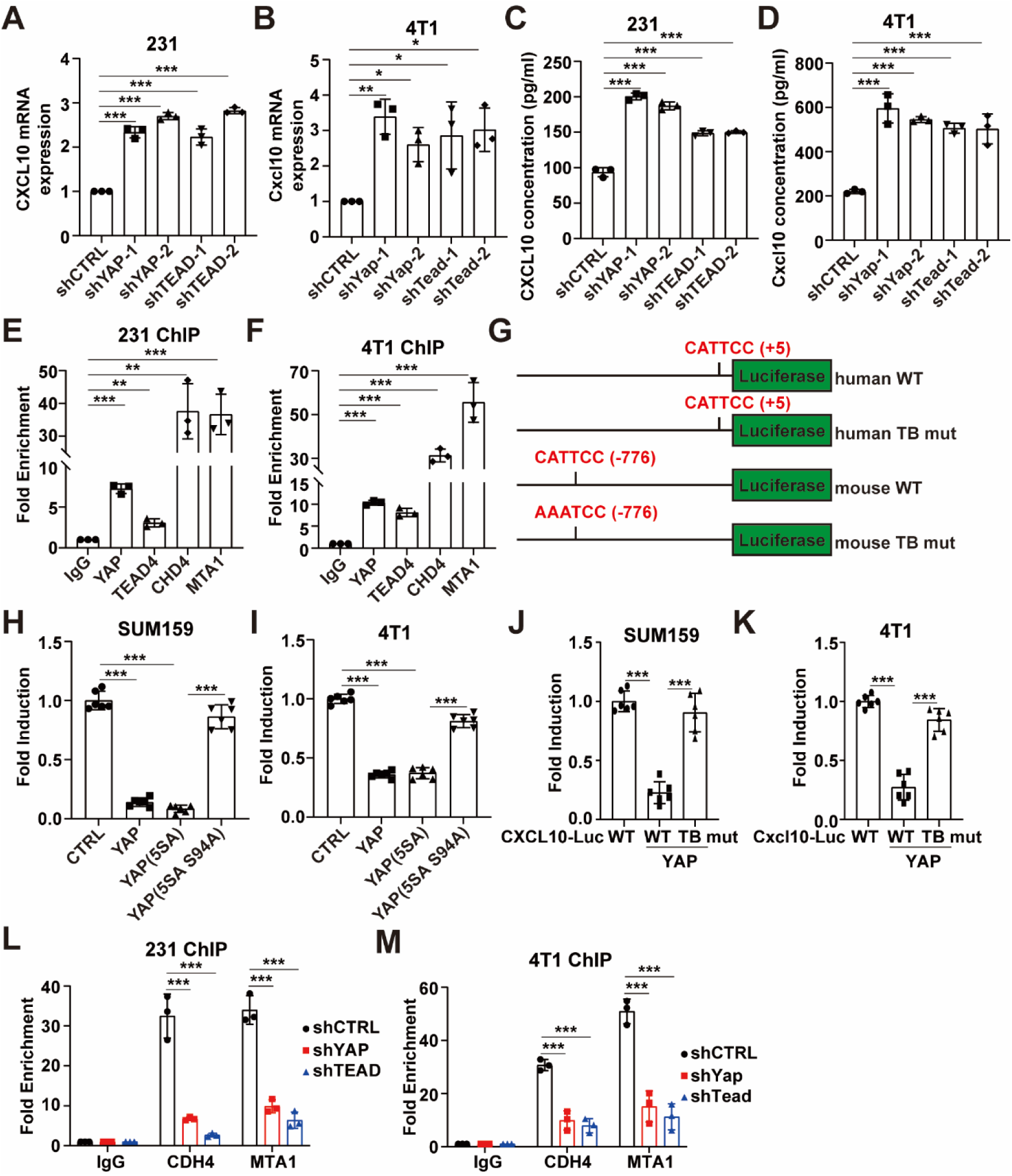
YAP/TEAD complex directly bind to CXCL10 promoter to repress its transcription. **(A)** qPCR analysis of the mRNA expression of CXCL10 in MDA-MB-231 cells with YAP or TEAD1/3/4 KD (n = 3). **(B)** qPCR analysis of the mRNA expression of Cxcl10 in 4T1 cells with Yap or Tead2/3/4 KD (n = 3). **(C)** ELISA analysis of CXCL10 protein expression in MDA-MB-231 cells with YAP or TEAD1/3/4 KD. **(D)** ELISA analysis of Cxcl10 protein expression in 4T1 cells with Yap or Tead2/3/4 KD. **(E)** ChIP-qPCR analysis of YAP, TEAD4, CHD4 and MTA1 signals using primers specific to CXCL10 promoter in MDA-MB- 231 cells. (n = 3). **(F)** ChIP-qPCR analysis of YAP, TEAD4, CHD4 and MTA1 signals using primers specific to Cxcl10 promoter in 4T1 cells (n = 3). **(G)** Schematic diagram showing the human and murine CXCL10 reporter constructs used in this study. The TEAD binding (TB) motifs and their positions with respect to the TSS are indicated in red. The numbers in parentheses refer to the positions of the first base of each TB motif. **(H)** Luciferase activity of human CXCL10 reporter in SUM159 cells transfected with the indicated expression plasmids (n = 6). **(I)** Luciferase activity of murine Cxcl10 reporter in 4T1 cells transfected with the indicated expression plasmids (n = 6). **(J)** Luciferase activity of human wild type or TB mutant CXCL10 reporter in SUM159 cells transfected with YAP (5SA) expression plasmids (n = 6). **(K)** Luciferase activity of murine wild type or TB mutant Cxcl10 reporter in 4T1 cells transfected with YAP (5SA) expression plasmids (n = 6). **(L)** ChIP-qPCR analysis of CHD4 and MTA1 signals using primers specific to CXCL10 promoter in MDA-MB-231 cells with YAP or TEAD1/3/4 KD (n = 3). **(M)** ChIP-qPCR analysis of CHD4 and MTA1 signals using primers specific to Cxcl10 promoter in 4T1 cells with YAP or TEAD2/3/4 KD (n = 3). Data are presented as means ± SD. *P* values in A, B C, D, E, F, H, I, J and K were determined by unpaired two-sided *t*-test. *P* values were calculated by one- way ANOVA in L and M. \**P* < 0.05, \*\**P* < 0.01, \*\*\**P* < 0.001.

Consistently, the mRNA level of YAP correlated negatively with CXCL10 mRNA in thyroid cancer and kidney cancer (fig. S7E).

To explore the molecular basis by which YAP or TEAD depletion enhanced CXCL10 expression, both human and mouse CXCL10 promoter sequences were analyzed by MatInspector (Genomatix) and observed harboring one putative TB sites (“CATTCC”) around the transcription start site (TSS) (fig. S5B). We also analyzed genomes of other species and found the CXCL10 TB motifs are conserved in primates, rodents and artiodactyla (fig. S5B). To confirm these predictions, we carried out ChIP-qPCR assays using anti-YAP or anti-TEAD4 antibody, and observed the occupancy of CXCL10 promoter near the TB sequence by YAP and TEAD4 in both human and mouse cells (Fig. 5, E and F).

To further elucidate whether CXCL10 is a directly repressed target gene of the YAP/TEAD complex, we separately cloned both human and mouse wild-type CXCL10 promoters (-1000 to +50 bp) and the according TB mutation promoters, and inserted them upstream of the luciferase gene (Fig. 5G). Overexpression of wild-type YAP or constitutively active YAP (5SA), but not TEAD-binding deficient YAP (5SA S94A) decreased the CXCL10 reporter activity, confirming the requirement of TEAD (Fig. 5, H and I). However, the TB motif mutation largely abrogated this repressive effect (Fig. 3, J and K).

Next, we investigated whether the NuRD complex involving in YAP/TEAD complex-mediated CXCL10 inhibition. Using ChIP-qPCR, both CHD4 and MTA1 were exhibited enrichment near the TB sequences of CXCL10 (Fig. 5, E and F), while the occupancy was reduced by YAP or TEAD KD (Fig. 5, L and M). On the basis of these results, we conclude that the YAP/TEAD complex recruits the NuRD complex to inhibit CXCL10 expression.

YAP loss enhances antitumor immunity via regulating NLRC5 expression Given YAP depletion induced NLRC5-mediated MHC-I APM expression and CTLs direct interaction with MHC-I on tumor cells, we hypothesized that YAP ablation or inhibition might facilitate CTLs mediated killing of tumor cells. Thus, to test this hypothesis, B16F10 cells expressing ovalbumin (OVA) (B16F10-OVA cells) were knocked down Yap and/or Nlrc5 or treated with YAP/TEAD inhibitor CA3. As expect, the presentation of OVA-derived peptide SIINFEKL by H-2Kb was enhanced by Yap KD or inhibition, while simultaneously depletion of Nlrc5 in Yap loss cells restored the presentation (fig. S8, A and B). Then using the OT-I transgenic mouse model (*28*), OT-I CD8^+^ T cells which engineered to express high affinity TCRs for OVA-derived peptide SIINFEKL were isolated and cocultured with the indicated B16F10-OVA cells (Fig. 6A and fig. S8C). Our results indicated that Yap-depletion or inhibited cells were significantly more efficient killing by CTLs than control cells (Fig. 6A and fig. S8C), and this phenomenon was largely repressed by simultaneously ablation of Nlrc5 in Yap loss cells (Fig. 6A).

**Fig. 6.**
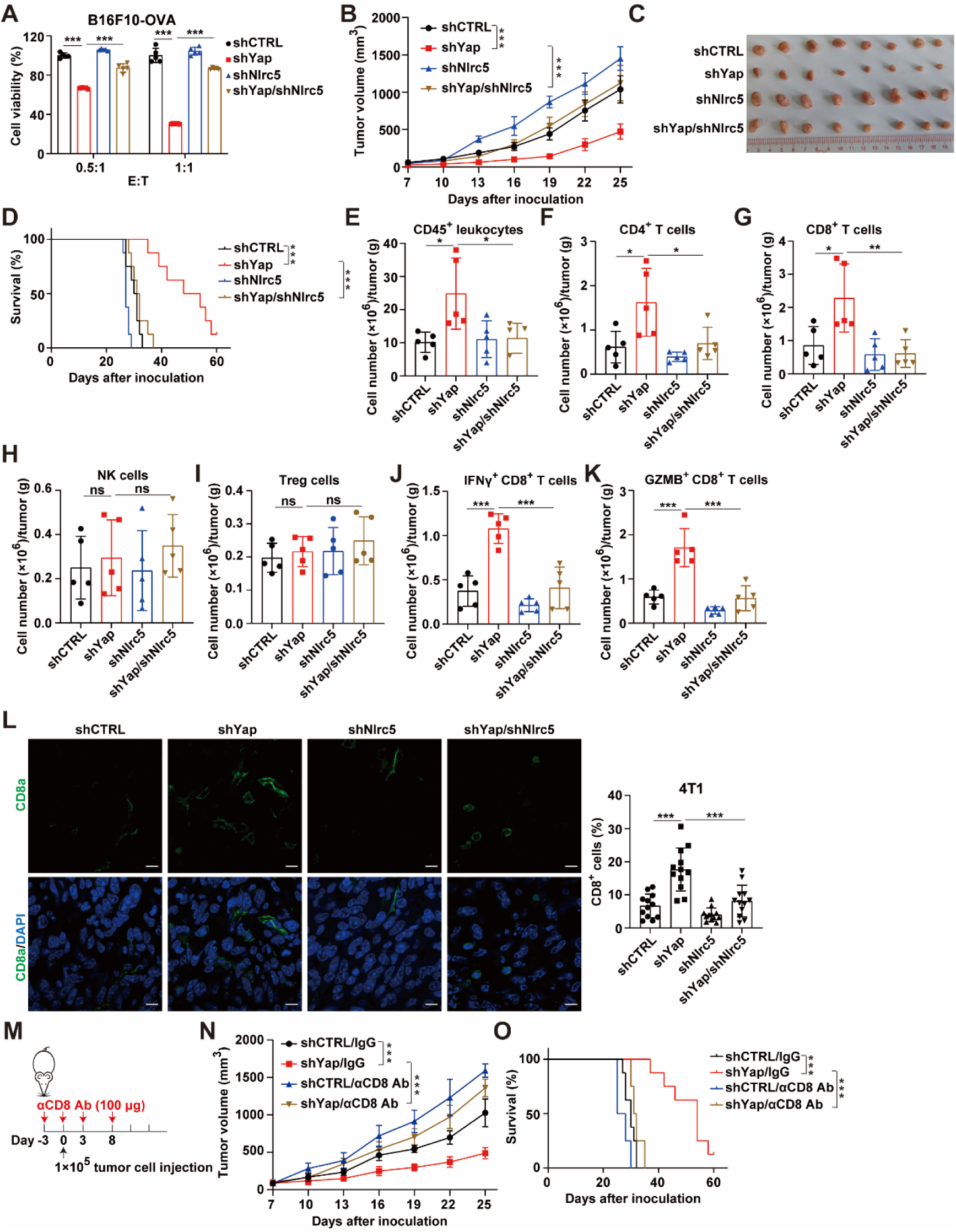
Depletion of YAP enhances CTL killing and intratumoral CTL infiltration dependent on NLRC5. **(A)** Coculturing of OT-1 CD8^+^ T cells with B16F10-OVA cells for 48 h and the B16F10-OVA cell viability was measured (n = 5). Effector to target (E : T) ratios are shown. **(B to D)** About 1 × 10^5^ mouse 4T1 tumor cells with Yap and/or Nlrc5 depletion were inoculated into the T4 mammary fat pad of BALB/c mice (n = 8). Tumor size **(B)**, tumor images **(C)** and overall survival **(D)** were monitored. **(E to I)** Flow Cytometric analysis of the indicated immune effector cells in 4T1 tumors (n = 5). **(J and K)** Flow Cytometric analysis of IFNγ^+^ and GZMB^+^ CD8^+^ T cells (n = 5). **(L)** Immunofluorescence staining (left) and quantitative estimates (right) of CD8a^+^ cells in 4T1 tumours (n = 3). Four fluorescent fields for each of the three samples were counted. Scale bar, 10 μm. **(M)** Injection schedule for antibody-mediated depletion of CD8^+^ T cells in BALB/c mice. **(N to O)** About 1 × 10^5^ mouse 4T1 tumor cells with or without Yap depletion were inoculated into the T4 mammary fat pad of BALB/c mice (n = 8). Tumor size **(N)** and overall survival **(O)** were monitored. Data are presented as means ± SD. *P* values in A, E, F, G, H, I, J, K, and L were determined by one-way ANOVA. *P* values were calculated by two-way ANOVA in B and N. *P* values in D and O were determined by long-rank test. \**P* < 0.05, \*\**P* < 0.01, \*\*\**P* < 0.001. ns, not significant.

We next evaluated the role of YAP loss induced NLRC5-mediated MHC-I APM expression in antitumor immunity. 4T1 or MC38 cells with Yap and/or Nlrc5 KD were inoculated into BALB/c and C57BL/6 mice, respectively. We found that simultaneously depletion of Nlrc5 largely reversed the inhibitory tumor growth and by Yap loss (Fig. 6, B to D, and fig. S8, D to F). To explore the tumor microenvirionment, the immune effector cells in tumors were quantified by flow cytometry (gating strategy in fig. S8S) and immunofluorescence staining. The flow cytometry analysis indicated that Yap ablation induced a profound increase in infiltration of CD45^+^ leukocytes (Fig. 6E and fig. S8G), CD4^+^ T helper cells (Fig. 6F and fig. S8H) and CD8^+^ T cells (CTLs) (Fig. 6G and fig. S8I), whereas no significant increases in NK cells (Fig. 6H and fig. S8J) and Treg cells (Fig. 6I and fig. S8K). Moreover, the numbers of interferon γ^+^ (IFNγ^+^) and granzyme B^+^ (GZMB^+^) CTLs also significantly increased after Yap KD (Fig. 6, J and K, and fig. S8, L and M). However, simultaneously depletion of Nlrc5 in Yap KD tumors restored the intratumoral infiltrating phenomenon of the immune effector cells (Fig. 6, E to K, and fig. S8, G to M). Immunofluorescence staining confirmed a significant increase in the number of intratumoral infiltrating CTLs (Fig. 6L and fig. S8N).

Consistent with our findings in mouse model, the mRNA level of YAP correlated negatively with the expression of CD8A in five human malignancies (fig. S8O).

To determine the role of CTLs, we depleted CTLs using anti-CD8 antibody, and found CTLs ablation completely reversed the growth delay of Yap loss tumors (Fig. 6, M to O, and fig. S8, P to R).

### YAP inhibition enhances ICB in a murine breast cancer model

Our results showing YAP depletion selectively stimulated MHC-I APM and CXCL10 expression, promoting us to evaluate whether YAP deficiency could synergize with ICB to enhance antitumor immunity. Control and Yap KD 4T1 cells were inoculated into the fourth left mammary glands of immunocompetent BALB/c mice. Then tumor-growth-delay studies were performed by a murine IgG control or anti-PD1 antibody (Fig. 7A). Compared with mice implanted with control cells, the tumors with Yap depletion exhibited increased sensitivity to anti-PD1 antibody (Fig. 7, B and C). Next, we extended the analysis using YAP inhibitor CA3 and anti-PD1 antibody (Fig. 7D). Although CA3 alone blocked tumor growth, the efficacy was significantly enhanced when combined with anti-PD1 antibody (Fig. 7, E and F). Accordingly, the intratumoral infiltrations of CTLs as well as IFNγ^+^ and GZMB^+^ CTLs were increased in Yap depletion tumors (Fig. 7, G to I). These observations confirmed that YAP represses antitumor immunity.

**Fig. 7.**
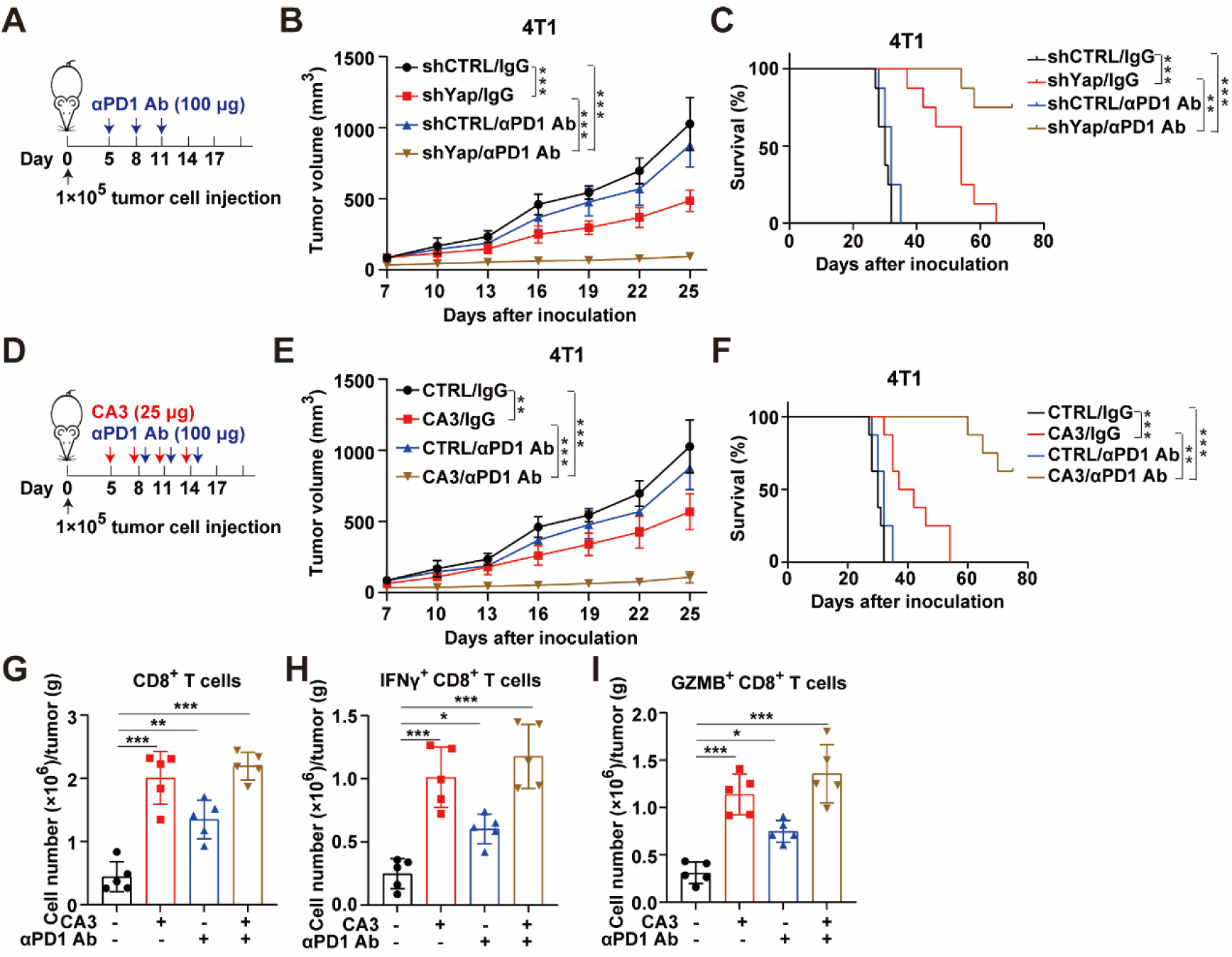
YAP inhibition overcomes tumor resistance to anti-PD1 ICB. **(A)** Treatment schedule for anti-PD1 antibody in BALB/c mice. **(B and C)** About 1 × 10^5^ mouse 4T1 tumor cells with or without Yap depletion were inoculated into the T4 mammary fat pad of BALB/c mice (n = 8). Tumor size **(B)**, and overall survival **(C)** were monitored. **(D)** Treatment schedule for anti-PD1 antibody combination with CA3 in BALB/c mice. **(E and F)** About 1 × 10^5^ mouse 4T1 tumor cells with or without Yap depletion were inoculated into the T4 mammary fat pad of BALB/c mice (n = 8). Tumor size **(E)**, and overall survival **(F)** were monitored. **(G)** Flow Cytometric analysis of CD8^+^ T cells (n = 5). **(H and I)** Flow Cytometric analysis of IFNγ^+^ and GZMB^+^ CD8^+^ T cells (n = 5). Data are presented as means ± SD. *P* values were calculated by two-way ANOVA in B and E. *P* values in C and F were determined by long-rank test. *P* values in G, H and I were determined by one-way ANOVA. \**P* < 0.05, \*\**P* < 0.01, \*\*\**P* < 0.001.

## Discussion

Immune evasion is still a major barrier for antitumor therapy. At present, only 10-30% of treated patients benefits from cancer immunotherapy (*16*). Dysfunction of cytotoxic CD8^+^ T lymphocytes (CTLs) caused by defect in antigen presentation has been implicated in immune evasion (*29*). Multiple studies have attempted to identify novel regulators of MHC-I and improve tumor antigen presentation (*13-18, 30, 31*). Here, our findings reveal cancer cell-intrinsic YAP activity antagonizes the antitumor immunity by regulating MHC-I antigen processing and presentation. We showed that YAP/TEAD complex directly binds and recruits NuRD complex to NLRC5 promoter to block NLRC5 transcription and NLRC5 mediated MHC-I APM gene expression, thereby suppress tumor-intrinsic immunogenicity and intratumoral infiltration of TCLs. Notably, YAP/TEAD inhibition sensitized the tumor to anti-PD1 ICB therapy. Taken together, our results provide a rationale of combination of YAP inhibitor and ICB therapy for cancer treatment.

The Hippo-YAP signaling is a crucial player in tumorigenesis and development. Dysregulated Hippo pathway and YAP/TEAD activity induces tumors in mouse models and occurs in various human carcinomas (*22*). Accumulating evidence has linked the Hippo-YAP signaling to the immune response in malignant neoplasms. The Hippo-YAP signaling has a vital role in the activity and functionality of T cells, B cells and macrophages, which is important for tumor immunity (*23, 24, 32-34*). In addition, the tumoral Hippo-YAP signaling directly affect the intratumoral infiltration and activation of T cells, MDSCs and macrophages and tumor-intrinsic PD-L1 expression (*35–40*). Most recently, Hippo pathway was identified as a novel regulator of cancer cell-intrinsic MHC-II expression (*41*). Our present study demonstrated that the Hippo-YAP signaling regulates the expression of MHC-I related genes in tumor cells, which proposes a novel immunomodulatory mechanism of the Hippo-YAP signaling.

Impaired antigen presentation caused by genetic mutation or epigenetic silencing of MHC-I related pathway is often associated with resistance to ICB therapy (*42, 43*). MHC-I transactivator (CITA) NLRC5 is an important coactivator of MHC-I related genes, and the expression of NLRC5 and MHC-I related genes is strongly induced in an IFN-γ- dependent manner (*19*). IFN-γ plays a two-sided function in antitumor immunity including potentiating antitumor immune response via enhancing MHC-I expression and also causing immune evasion through upregulating at least PD-L1 expression to (*44*). We showed here YAP loss promotes NLRC5 transcription and the expression of MHC-I related genes in the absence of IFN-γ. Thus, YAP may be uniquely positioned to be an immunogenic target for cancer immunotherapy.

In addition to regulate MHC-I expression, our results also confirmed the YAP deficiency induces chemokine CXCL10 expression (Fig. 5), which attracts CTLs recruitment into tumors (*45*). Considering this and TCR of TCLs directly interacting with MHC-I on cancer cells, we focus on CTLs. However, the changes in other immune cells such as CD4^+^ T cells after YAP KD were also observed. Given YAP inactivation promotes cGAS– STING signaling (*46*), further investigations are needed to elucidate how these changes in other immune cells are mediated.

In most circumstances, YAP is a potent transcriptional coactivator to upregulate target gene expression such as CTGF (*47*). However, YAP can also function as a transcriptional corepressor (*26*). The repressor activity of YAP was reported to be mediated by the NuRD complex (*48*). Our results proposed that the YAP/TEAD complex directly binds to NLRC5 promoter and guided the NuRD complex to the promoters, thereby repressing NLRC5 transcription.

In summary, our findings suggest that YAP as a cancer cell intrinsic regulator of cell-surface MHC-I related pathway and chemokine CXCL10 and therefore influences intratumoral immune infiltration, which proposes pharmaceutical inhibition of YAP may sensitize a subset of patients to ICB treatment.

## Materials and Methods

### Constructs

The full-length human NF2 or YAP cDNA was cloned into Flag-CMV2 vector. Human MST2 expression plasmid was generously provided by Kunliang Guan (University of California, La Jolla, CA). Human LATS1 expression plasmids were obtained from Addgene (41156). Human and murine NLRC5 and CXCL10 promoters were cloned into pGL3 basic luciferase reporter vector (Promega). All mutated plasmids were constructed from wild type plasmids by Fast Mutagenesis System (TransGen Biotech). The shRNAs were cloned into pLKO.1 vector. The sequences of these shRNAs are shown in table S1. All the cloned sequences were confirmed by sequencing.

### Cell culture and treatment

HEK293T, MDA-MB-231, SUM159, 4T1 and MC38 cell lines were obtained from American Type Culture Collection (ATCC) and the B16F10- OVA cells were kindly provided by Qiming Liang’s lab (Shanghai Jiao Tong University, China). HEK293T and MDA-MB-231 cells were maintained in DMEM supplemented with 10% fetal bovine serum (FBS) and 1% penicillin-streptomycin at 37 °C and 5% CO_2_. SUM159, 4T1 and MC38 cells were cultured in RPMI-1640 supplemented with 10% FBS and 1% penicillin-streptomycin at 37 °C and 5% CO_2_. All cells were tested for mycoplasma using the MycoBlue Mycoplasma Detector Kit (Vazyme). Transfection of plasmids or siRNA duplexes was performed with Lipofectamine 3000 (Thermo Fisher Scientific) according to the manufacturer’s instructions. Stable cell lines were established with a lentiviral infection followed by selection with an appropriate antibiotic. Hydrogel assay were performed as previously described (*49*).

### CD8^+^ T-cell cytotoxicity assay

Mouse CD8^+^ T cells were isolated from the spleens of OT-I mice using the EasySep^TM^ Mouse CD8^+^ T Cell Isolation Kit (STEMCELL, 19853) following the manufacturer’s instructions and then cultured in complete RPMI-1640 media (10% FBS, 20mM HEPES, 2 mM L-glutamine, 1 mM sodium pyruvate, 1% penicillin-streptomycin) supplemented with 50 mM 2-mercaptoethanol and 1 μg/ml OVA_257-264_ (SIINFEKL) peptide (GenScript) at at 37 °C and 5% CO_2_ for 3 days. After removing the peptide, CD8^+^ T cells were incubated with 30 U/ml mouse recombinant interleukin- 2 (IL-2) for an additional 2 days. To tested the sensitivity of cancer cells to T cell-driven cytotoxicity, ten thousand B16F10-OVA cells and five or ten thousand CD8^+^ T cells were cocultured in complete RPMI-1640 media in each well of 96-well plates for 48 h. Then the viability of B16F10-OVA cells was measured by the Lactate Dehydrogenase A Release Assay Kit (Beyotime).

### Western blotting

Cells were lysed in RIPA buffer (50 mM Tris-HCl, pH7.4, 150 mM NaCl, 1% NP-40, 0.1% SDS) supplemented with phosphatase-inhibitor cocktail, protease-inhibitor cocktail and 1 mM PMSF followed by mild sonication. Equal proteins were separated by SDS-PAGE and transferred to PVDF membranes. Western blotting was performed with the indicated primary antibodies (NLRC5 (Abclonal, A16740), β2-microglobulin (Cell Signaling Technology, 12851S), TAP1 (Cell Signaling Technology, 12341S), YAP (Cell Signaling Technology, 14074S), p-YAP (Cell Signaling Technology, 4911S), MHC-I (Santa Cruz, sc-32235), PSMB9 (Abcam, ab242061), TEAD4 (Abnova, H00007004-M01), Flag (Sigma-Aldrich, F1804), Myc-Tag (Abmart, M200022), Actin (Proteintech, 66009-1-1g)), and then with the related horseradish peroxidase conjugated secondary antibody. The membrane was then detected with ECL Plus Western Blotting Substrate (Millipore) using Chemiluminescent Imaging System (Tanon 5200, China). RNA isolation and real-time PCR assay Total RNA was extacted by RNAiso Plus (TaKaRa) according to the manufacturer’s protocol. First strand cDNA was synthesized using the Primescript RT Reagent Kit (TaKaRa) and 0.5 μg of total RNA in each cDNA synthesis reaction. Then prepared cDNA was subjected to real-time PCR analysis with Eastep qPCR Master Mix (Promega) and primer mixtures (primer sequences are shown in table S2). The relative expression of genes was normalized against GAPDH and calculated by the comparative Ct method.

### ChIP-qPCR assay

The ChIP assay was carried out using the EZ-Magna ChIP A/G Chromatin Immunoprecipitation Kit (Millipore) following the manufacturer’s instructions. YAP (Cell Signaling Technology, 14074S), TEAD4 (Abnova, H00007004-M01), CHD4 (Abcam, ab70469), and MTA1 (Cell Signaling Technology, 5646S) antibodies were used for immunoprecipitation. Then the eluted DNA was subjected to real-time PCR analysis with Eastep qPCR Master Mix (Promega) and promoter-specific primer mixtures. The primer sequences are listed in table S2.

### Lentivirus production and infection

Lentiviral packaging was performed using HEK293T cells. Transfection was performed using Polyethylenimine (PEI) (Polysciences, 23966) according to the manufacturer’s protocol. For each 10-cm dish, 10 μg of pLKO.1 vector, 2.5 μg of pMD2.G and 7.5 μg of psPAX2 were used. Then fifty microliters of PEI (1 mg/ml) were diluted by 1.5 ml Opti-MEM and gently added into the DNA mixture. After incubation for 15-20 minutes, the transfection mix was then carefully added to 293T cells. The culture supernatants containing the lentivirus were collected at 48 h and 72 h after transfection. For infection with lentivirus, cells were cultured with lentiviral solution for 8-12 h in the presence of 8 μg/mL Polybrene (Sigma-Aldrich).

### Immunofluorescence

For immunofluorescence analyses, tumors from mice were fixed in 4% formaldehyde in PBS for 24 h, then dehydrated and embedded into cryo-embedding media (OCT), sectioned at 8 μm and then mounted onto slides.

Then the immunostaining was performed by standard procedures using primary antibody against mouse CD8a (Abcam, ab217344) and then with the related fluorescent-labelled secondary antibody. For visualization of cell nucleus, DAPI was used. After washing with PBS, the stained slides were mounted with mounting medium. Imaged were captured using a fluorescence microscope (LSM880, ZEISS).

### Luciferase reporter assay

Cells in 24-well plates were transfected with 250 ng of reporter plasmid, 50 ng of control plasmid pCMXβgal, and 200 ng of the indicated expression plasmids using Lipofectamine 3000 according to the manufacturer’s protocol. After transfection for 48 h, the luciferase activity was measured using a luciferase reporter assay kit (Promega, E1501) and the luciferase values were normalized according to β-gal activity.

### RNA-seq and data analysis

For comparison of transcription profile, total RNA was extracted from control and YAP KD 4T1 cells, and then submitted to Illumina HiSeq™ 2500, which was performed by Gene Denovo Biotechnology Company (Guangzhou, China). Sequencing reads were filtered by fastp (version 0.18.0), mapped to the reference genome using HISAT2(version 2.1.0), and then assembled by Stringtie (version 1.3.4). A FPKM (fragment per kilobase of transcript per million mapped reads) value was calculated to quantify the expression abundance and variations of each transcription region by RSEM software. Statistics for differentially expressed genes (DEGs) with a false discovery rate (FDR) below 0.05 and absolute fold change ≥ 2.0 were performed by edgeR. Enrichment analyses of all DEGs were carried out by the Gene Ontology (GO) database and the Kyoto Encyclopedia of Genes and Genomes (KEGG) database using p < 0.05 as the threshold.

### Flow cytometry assay

For in vitro cell-surface staining, cells were harvested and incubated with the indicated antibodies at 4 °C for 30 min. For the immunophenotyping, tumors were collected, weighed, chopped with scissors and incubated in RPMI-1640 containing 1 mg/ml collagenase IV (Worthington), 100 μg/ml DNase I (Sigma-Aldrich), 1% FBS for 40 min at 37 °C. The dissociated cells were passed through a 70-μm tube-secured cell strainer (Falcon). Then one million cells were blocked with an anti-CD16/32 antibody (BioLegend) for 10 min at 4 °C followed by staining with the following appropriate antibodies for 30 min at 4 °C. For intracellular cytokine staining, cells were stimulated with PMA (10 ng/ml), ionomycin (500 ng/ml) and BFA (1 μg/ml) for 8 h at 37 °C followed by staining with a fixation/permeabilization buffer according to the manufacturer’s protocol (BioLegend). After washing, the analyses were performed by a Beckman CytoFlex flow cytometer. Gating strategies were shown in fig. S8. The following antibodies were used from BioLegend: PE anti-mouse β2-microglobulin (154505, clone A16041A); APC anti-mouse MHC-I (H- 2Kd/H-2Dd) (114714, clone 34-1-2S); APC anti-mouse MHC-I (H-2Kb) (116518, clone AF6-88.5); APC anti–mouse H-2Kb: SIINFEKL (141606, clone 25-D1.16); APC anti–human HLA-A/B/C (311410, clone W6/32); FITC anti-human β2-microglobulin (316304, clone 2M2); APC anti-mouse CD45.2 (109814, clone 104); Brilliant Violet 421 anti-mouse CD3 (100341, clone 145-2C11); FITC mouse anti-CD3 (100305, clone 145-2C11); PerCP-cyanine 5.5 anti-mouse CD4 (100434, clone GK1.5), FITC anti-mouse CD8a (100706, clone 53-6.7); PE anti-mouse NK-1.1 (108706, clone PK136); PE anti-mouse Foxp3 (126403, clone MF-14); PE anti-mouse IFN-γ (505808, clone XMG1.2); PE/Cyanine7 anti-Granzyme B (372214, clone QA16A02).

### Animal model studies

All animal experimental protocols were approved by the Animal Research Ethics Committee of Shenzhen University (permit number: AEWC- 201412003). NOD/ShiLtJGpt-Prkdcem26Cd52Il2rgem26Cd22/Gpt (NCG) mice, BALB/c mice and C57BL/6 mice were purchased from GemPharmatech Company (Nanjing, China). OT-I transgenic mice in the C57BL/6 background were obtained from The Jackson Laboratory (Bar Harbor, ME). All the mice were housed in a specific pathogen free facility with six mice per cage at constant temperature and humidity under a light cycle of 12 h on/12 h off set from 6 am to 6 pm. Mice were randomly divided and at least five mice per treatment group were included. For 4T1 mammary carcinoma models, 1 × 10^5^ 4T1 cells were implanted into the T4 mammary fat pad of 8-week-old female NCG or BALB/c mice. For MC38 colon adenocarcinoma models, every 8-week-old male C57BL/6 mouse was subcutaneously (s.c.) inoculated with 1 × 10^5^ MC38 cells. To depletion of CD8^+^ T cells, mice were intraperitoneally (i.p.) injected with 200 μg isotype control (BioXCell, BE0088) or neutralizing antibody against CD8 (clone 53-5.8, BioXCell, BE0223) on days -3, 0, 3 and 8. For ICB, mice were i.p. injected with 100 μg isotype control (BioXCell, BE0089) or anti- PD1 antibody (clone RMP1-14, BioXCell, BE0146) on days 5, 8 and 11. To test the combination efficacy of YAP inhibitor CA3 and ICB, starting on day 5 after transplantation, 25 μg CA3 (Selleck, S8661) or equal vehicle (5% DMSO/40% PEG 300/5% Tween 80 in saline) was i.p. injected, then both 25 μg CA3 or equal vehicle and 100 μg isotype control or anti- PD1 antibody were i.p. administrated on days 8, 11 and 14. Tumor sizes were measured every three days with a caliper and tumor volumes were calculated using the formula: 0.5 × length × width^2^. To assess the end point of animals, mice bearing tumors reaching 1.5 cm in its longest dimension or 2,000 mm^3^ in diameter were defined as dead.

### Cytokine measurement

The ELISA analyses were performed to measure CXCL10 levels in human and mouse cell culture supernatants by Human IP-10/CXCL10 ELISA Kit (QuantiCyto, EHC157) and Mouse IP-10/CXCL10 ELISA Kit (QuantiCyto, EMC121) according to the manufacturer’s instructions, respectively.

### Statistical analysis

Statistical analysis was carried out with GraphPad Prism 9 software. The data were analyzed by unpaired two-sided Student’s t test, One-way ANOVA, Two-way ANOVA or Long-rank test. Results are presented as mean ± SD. We used the Gene Expression across Normal and Tumour tissue database (GENT) (*50*) to analyze the relationship between YAP and NLRC5, HLA-A, B2M, PSMB9, TAP1, CXCL10, CD8A in indicated patient cohorts. For all the analyses, statistical significance was set to *P* < 0.05.

## Acknowledgments

We thank Qiming Liang at Shanghai Jiao Tong University, China for providing B16F10-OVA cells, Instrument Analysis Center of Shenzhen University for the assistance with immunofluorescence analysis, and the Cellular and Molecular Biology Center of China Pharmaceutical University for the assistance with FACS analysis.

## Author contributions

ZW, MQ and DL developed the concept and designed this work. LP, LZ, HL, XZ, KW, XM, DZ, SX, YD and TW performed the experiments and carried out the data acquisition. HL, ZW, MQ and DL performed data analysis. ZW, MQ and DL edited and revised the manuscript. All authors read and approved this manuscript.

---

**Fig. S1.**
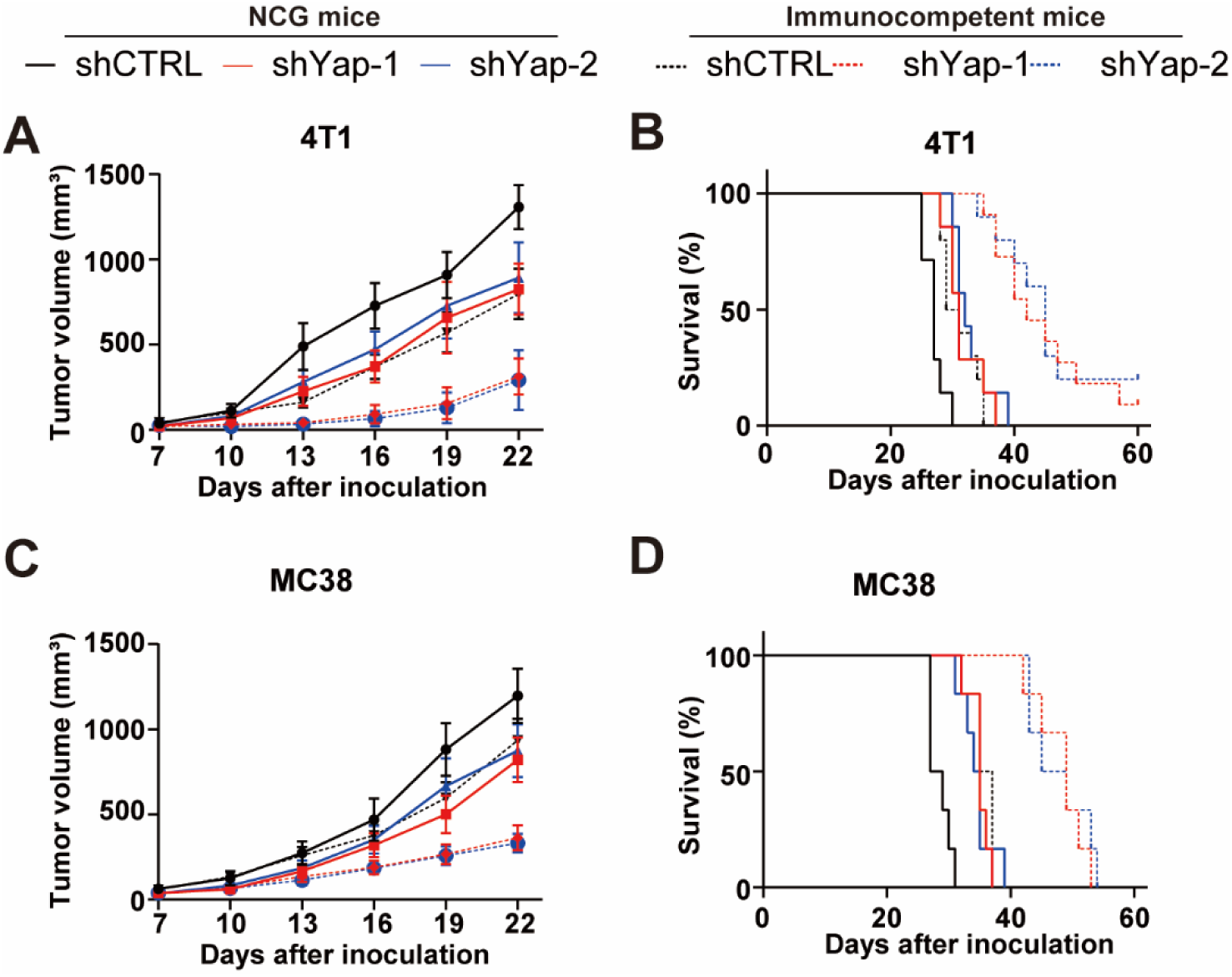
YAP ablation induces a more obvious reduction of tumor growth in syngeneic immunocompetent mice than severe immunodeficient NCG mice. (A and B) About 1 × 10^5^ mouse 4T1 tumor cells with or without Yap depletion were inoculated into the T4 mammary fat pad of NCG mice (n = 7) or BALB/c mice (n = 9). Both tumor size **(A)** and overall survival **(B)** were monitored. **(C and D)** About 1 × 10^5^ mouse MC38 tumor cells with or without Yap depletion were subcutaneously implanted into NCG mice (n = 6) or BALB/c mice (n = 6). Both tumor size **(C)** and overall survival **(D)** were monitored. Data are shown as means ± SD.

**Fig. S2.**
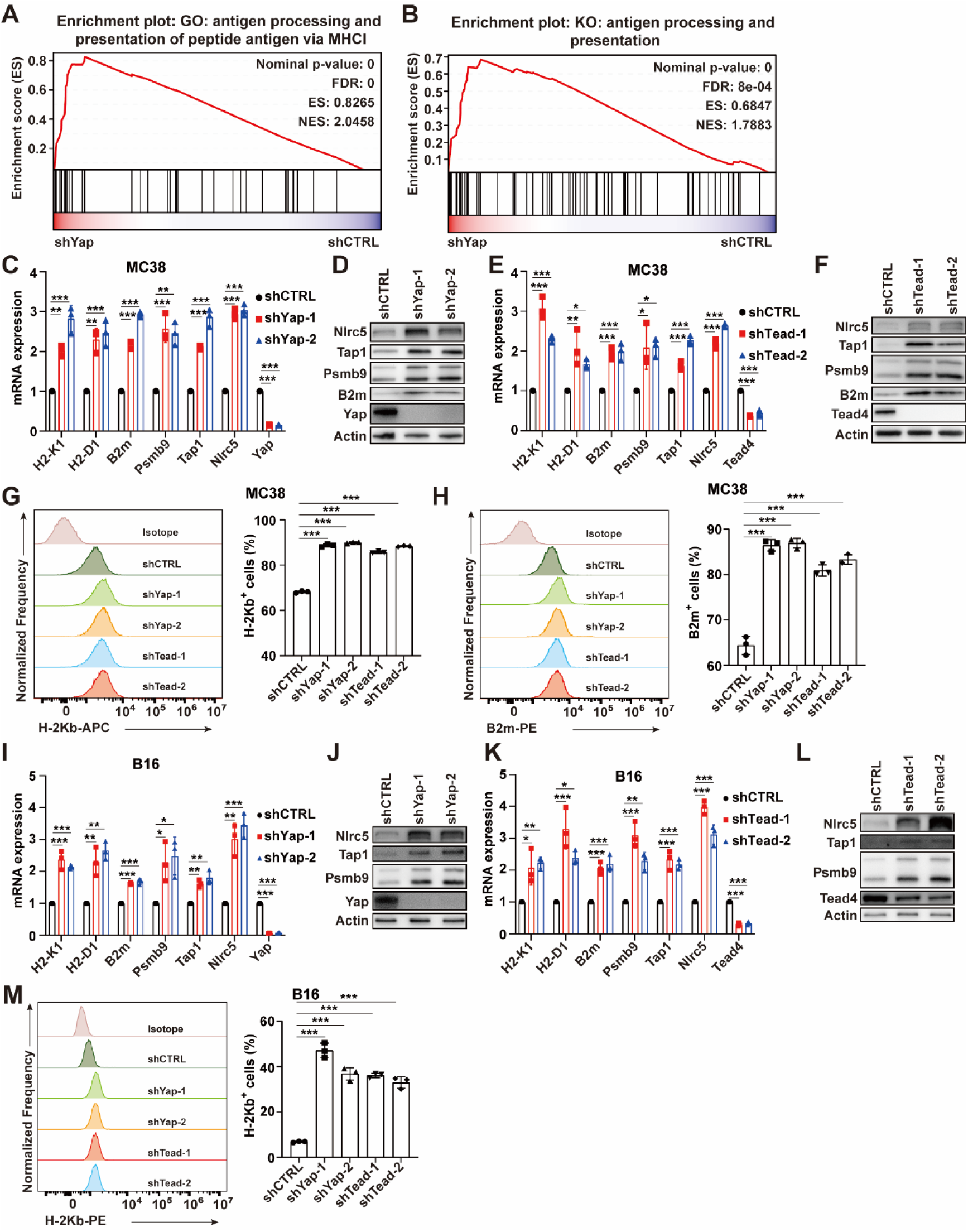
YAP or TEAD knockdown increases the expression of NLRC5 and MHC-I APM genes in mouse tumor cells. **(A)** Gene set enrichment analysis (GSEA) of Gene Ontology (GO) pathways using genes differentially expressed between 4T1 cells with or without Yap knockdown (KD). **(B)** Gene set enrichment analysis (GSEA) of KEGG pathways using genes differentially expressed between 4T1 cells with or without Yap KD. (**C**) qPCR analysis of the mRNA expression of MHC-I APM genes and Nlrc5 gene in MC38 cells with or without Yap KD (n = 3). **(D)** Immunoblot (IB) analysis of MC38 cells with or without Yap KD. **(E)** qPCR analysis of the mRNA expression of MHC-I APM genes and Nlrc5 gene in MC38 cells with or without Tead2/3/4 KD (n = 3). **(F)** IB analysis of MC38 cells with or without Tead2/3/4 KD. **(G)** Flow cytometry detection of cell-surface level of H-2Kb in MC38 cells with or without Yap or Tead2/3/4 KD. The percentage of H-2Kb positive cells is shown (n = 3). **(H)** Flow cytometry detection of cell-surface level of B2m in MC38 cells with or without Yap or Tead2/3/4 KD. The percentage of B2m positive cells is shown (n = 3). Data are presented as means ± SD. *P* values were calculated by one-way ANOVA in C and E. *P* values in G and H were determined by unpaired two-sided *t*-test. \**P* < 0.05, \*\**P* < 0.01, \*\*\**P* < 0.001.

**Fig. S3.**
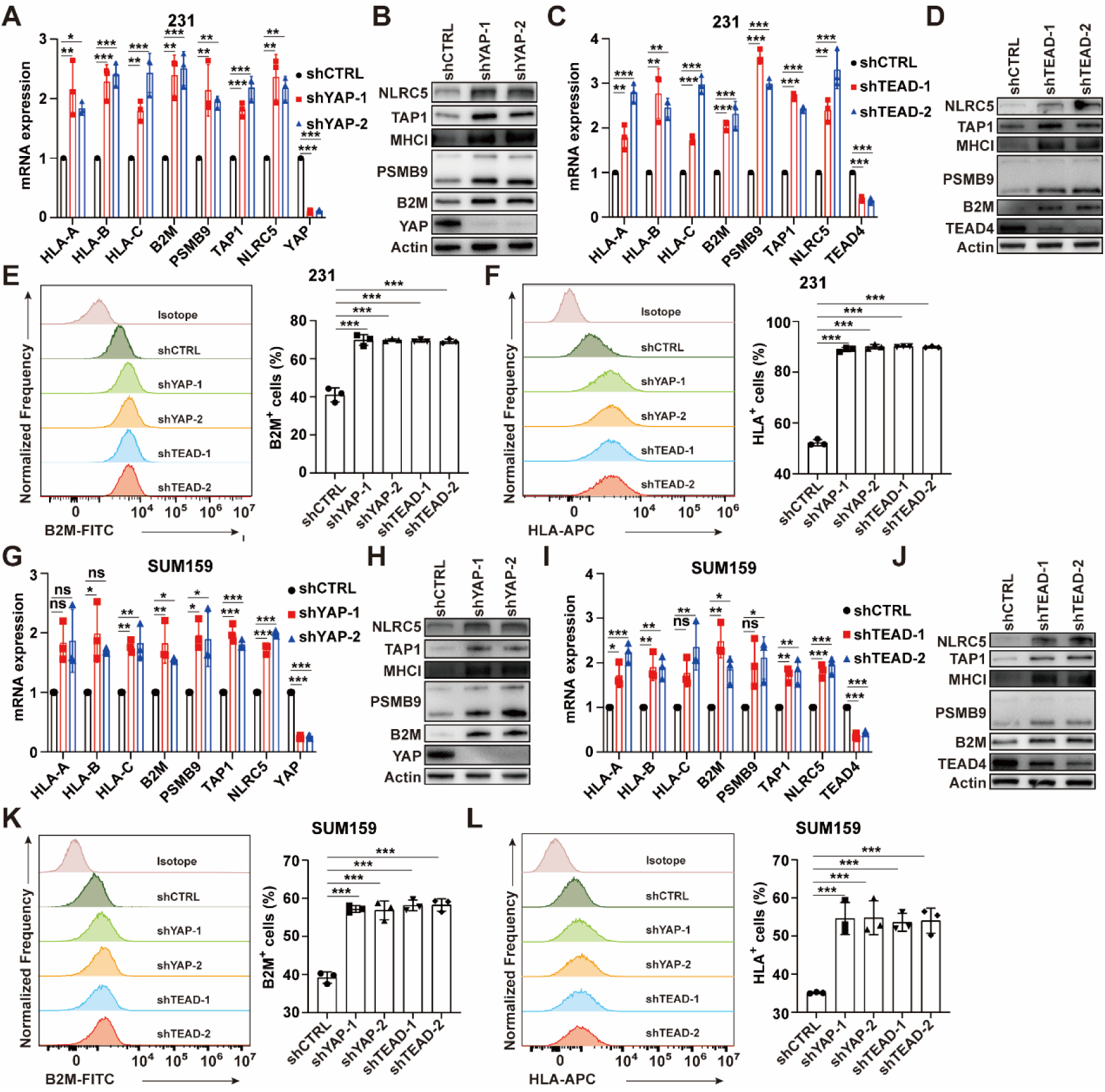
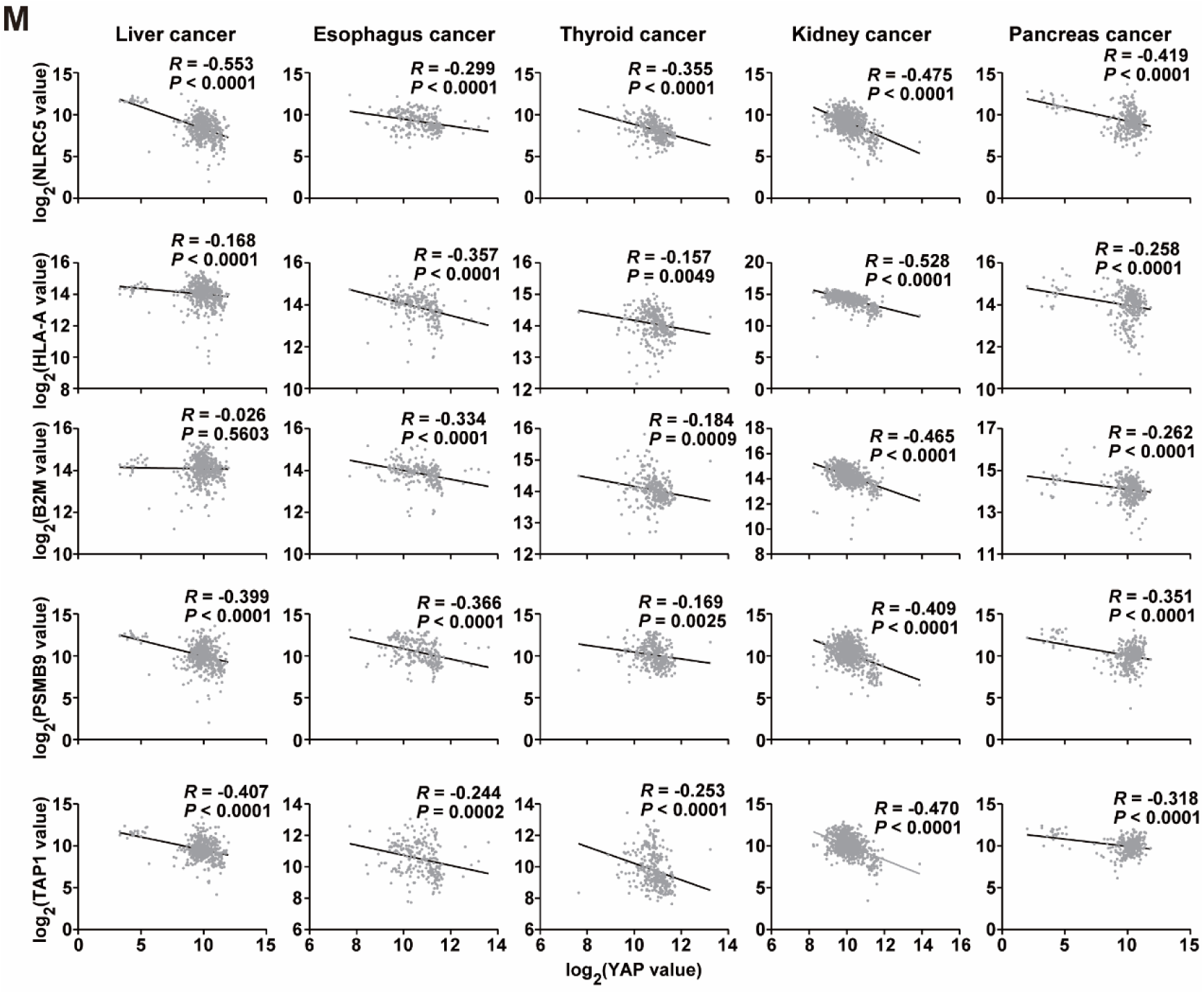
YAP or TEAD depletion ehhances the expression of NLRC5 and MHC-I APM genes in human tumor cells. **(A)** qPCR analysis of the mRNA expression of MHC-I APM genes and NLRC5 gene in MDA-MB- 231 cells with or without YAP KD (n = 3). **(B)** Immunoblot (IB) analysis of MDA-MB-231 cells with or without YAP KD. **(C)** qPCR analysis of the mRNA expression of MHC-I APM genes and NLRC5 gene in MDA-MB- 231 cells with or without TEAD1/3/4 KD (n = 3). **(D)** IB analysis of MDA-MB-231 cells with or without TEAD1/3/4 KD. **(E)** Flow cytometry detection of cell-surface level of HLA-A/B/C in MDA-MB-231 cells with or without YAP or TEAD1/3/4 KD. The percentage of HLA-A/B/C positive cells is shown (n = 3). **(F)** Flow cytometry detection of cell-surface level of B2M in MDA-MB-231 cells with or without YAP or TEAD1/3/4 KD. The percentage of B2M positive cells is shown (n = 3). **(G)** qPCR analysis of the mRNA expression of MHC-I APM genes and NLRC5 gene in SUM159 cells with or without YAP KD (n = 3). **(H)** Immunoblot (IB) analysis of SUM159 cells with or without YAP KD. **(I)** qPCR analysis of the mRNA expression of MHC-I APM genes and NLRC5 gene in SUM159 cells with or without TEAD1/3/4 KD (n = 3). **(J)** IB analysis of SUM159 cells with or without TEAD1/3/4 KD. **(K)** Flow cytometry detection of cell-surface level of HLA-A/B/C in SUM159 cells with or without YAP or TEAD1/3/4 KD. The percentage of HLA-A/B/C positive cells is shown (n = 3). **(L)** Flow cytometry detection of cell-surface level of B2M in SUM159 cells with or without YAP or TEAD1/3/4 KD. The percentage of B2M positive cells is shown (n = 3). **(M)** Negative correlation of YAP mRNA levels with that of NLRC5 and MHC-I APM genes in human liver cancer (517 samples), esophagus cancer (236 samples), thyroid cancer (320 samples), kidney cancer (810 samples) and pancreas cancer (324 samples). Data from the Gene Expression across Normal and Tumor tissue (GENT) database. *R* represents the Pearson correlation coefficient. Data are presented as means ± SD. *P* values were calculated by one-way ANOVA in A, C, G and H. *P* values in E, F, K and L were determined by unpaired two-sided *t*-test. *P* value in M was calculated by *F*-test. \**P* < 0.05, \*\**P* < 0.01, \*\*\**P* < 0.001.

**Fig. S4.**
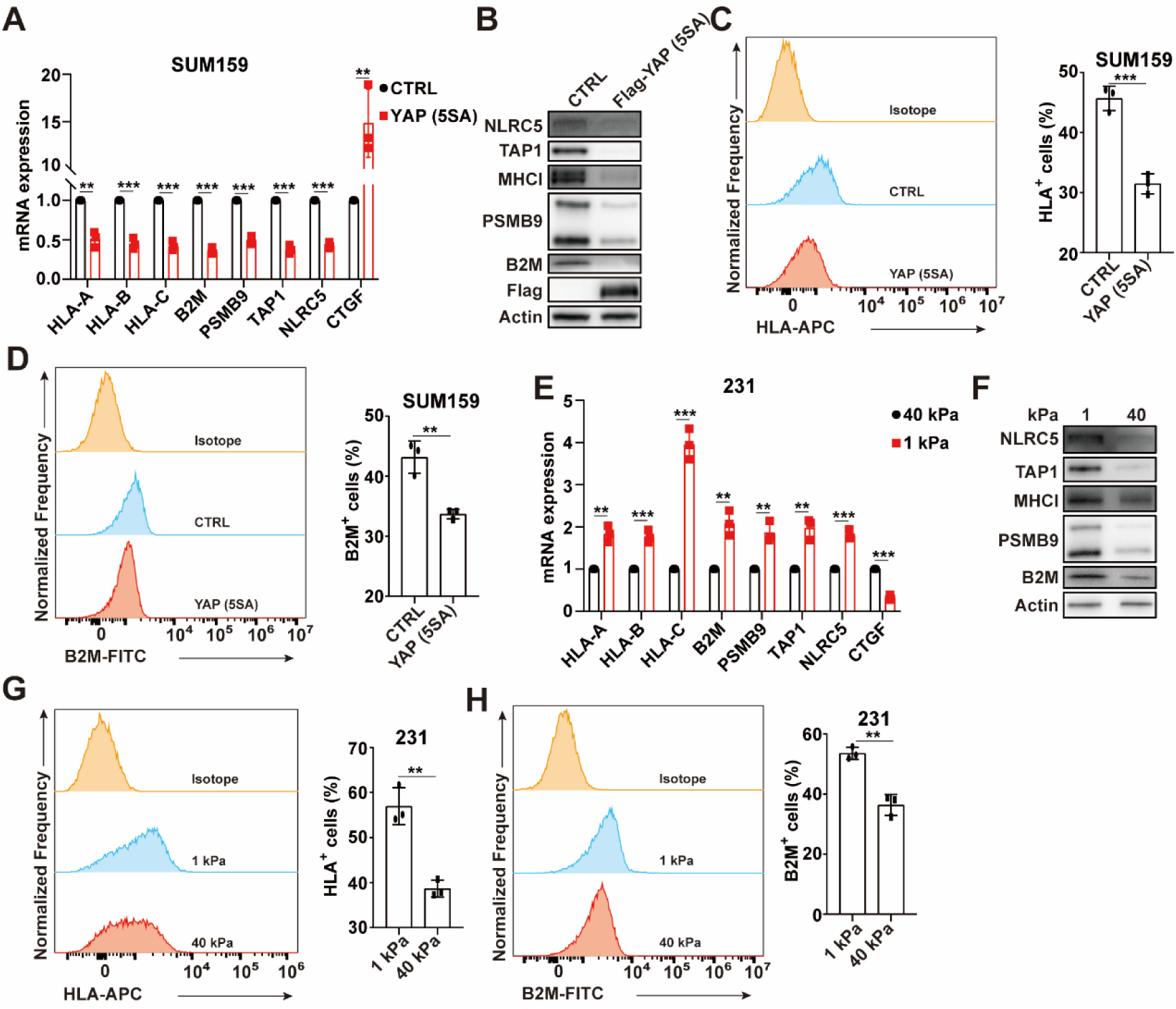
The Hippo pathway modulates NLRC5 and MHC-I APM gene expression. **(A)** qPCR analysis of the mRNA expression of MHC-I APM genes and NLRC5 gene in SUM159 cells transfected with or without constitutive active YAP (5SA). A Hippo target gene CTGF is used as positive control (n = 3). **(B)** Immunoblot (IB) analysis of SUM159 cells transfected with or without YAP (5SA). **(C)** Flow cytometry detection of cell-surface level of HLA-A/B/C in SUM159 cells transfected with or without YAP (5SA). The percentage of HLA-A/B/C positive cells is shown (n = 3). **(D)** Flow cytometry detection of cell-surface level of B2M in SUM159 cells transfected with or without YAP (5SA). The percentage of B2M positive cells is shown (n = 3). **(E)** qPCR analysis of the mRNA expression of MHC-I APM genes and NLRC5 gene in MDA-MB-231 cells grown on stiff (elastic modulus of 40 kPa) or soft (1 kPa) fibronectin-coated hydrogels. CTGF is used as positive control (n = 3). **(F)** IB analysis of MDA-MB-231 cells grown on stiff (elastic modulus of 40 kPa) or soft (1 kPa) fibronectin-coated hydrogels. **(G)** Flow cytometry detection of cell-surface level of HLA-A/B/C in MDA-MB-231 cells grown on stiff (elastic modulus of 40 kPa) or soft (1.0 kPa) fibronectin-coated hydrogels. The percentage of HLA-A/B/C positive cells is shown (n = 3). **(H)** Flow cytometry detection of cell-surface level of B2M in MDA-MB-231 cells grown on stiff (elastic modulus of 40 kPa) or soft (1.0 kPa) fibronectin-coated hydrogels. The percentage of B2M positive cells is shown (n = 3). Data are presented as means ± SD. *P* values were determined by unpaired two-sided *t*-test. \*\**P* < 0.01, \*\*\**P* < 0.001.

**Fig. S5.**
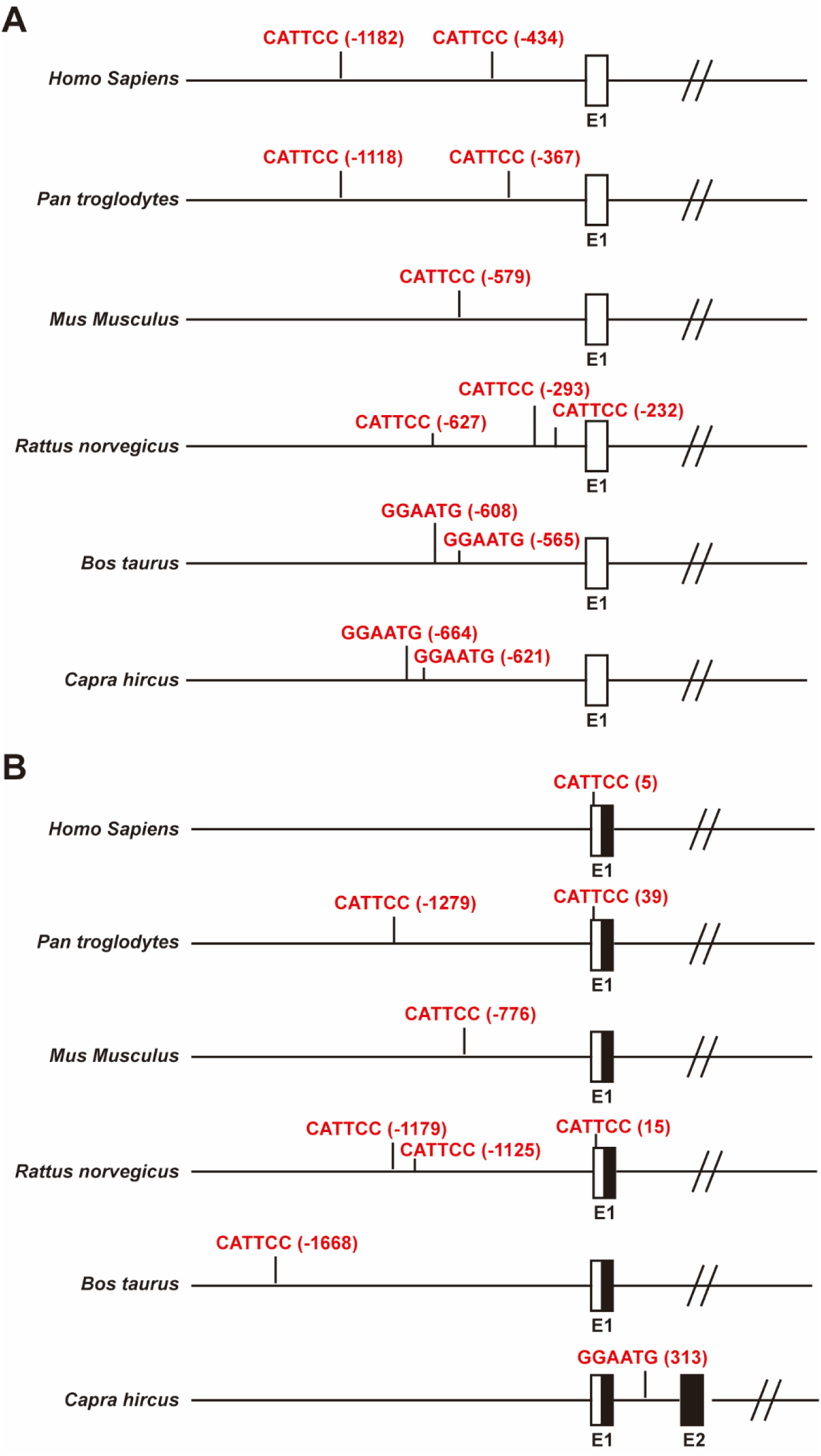
TEAD binding (TB) motifs in the NLRC5 and CXCL10 genes are evolutionarily conserved. Schematic diagram showing the TSS regions of the NLRC5 **(A)** and CXCL10 **(B)** loci in the indicated species. The TB motifs and their positions with respect to the TSS are indicated in red. The numbers in parentheses refer to the positions of the first base of each TB motif. Transcription proceeds from left to right. Boxes indicate exons, with black indicating coding regions and white indicating untranslated regions.

**Fig. S6.**
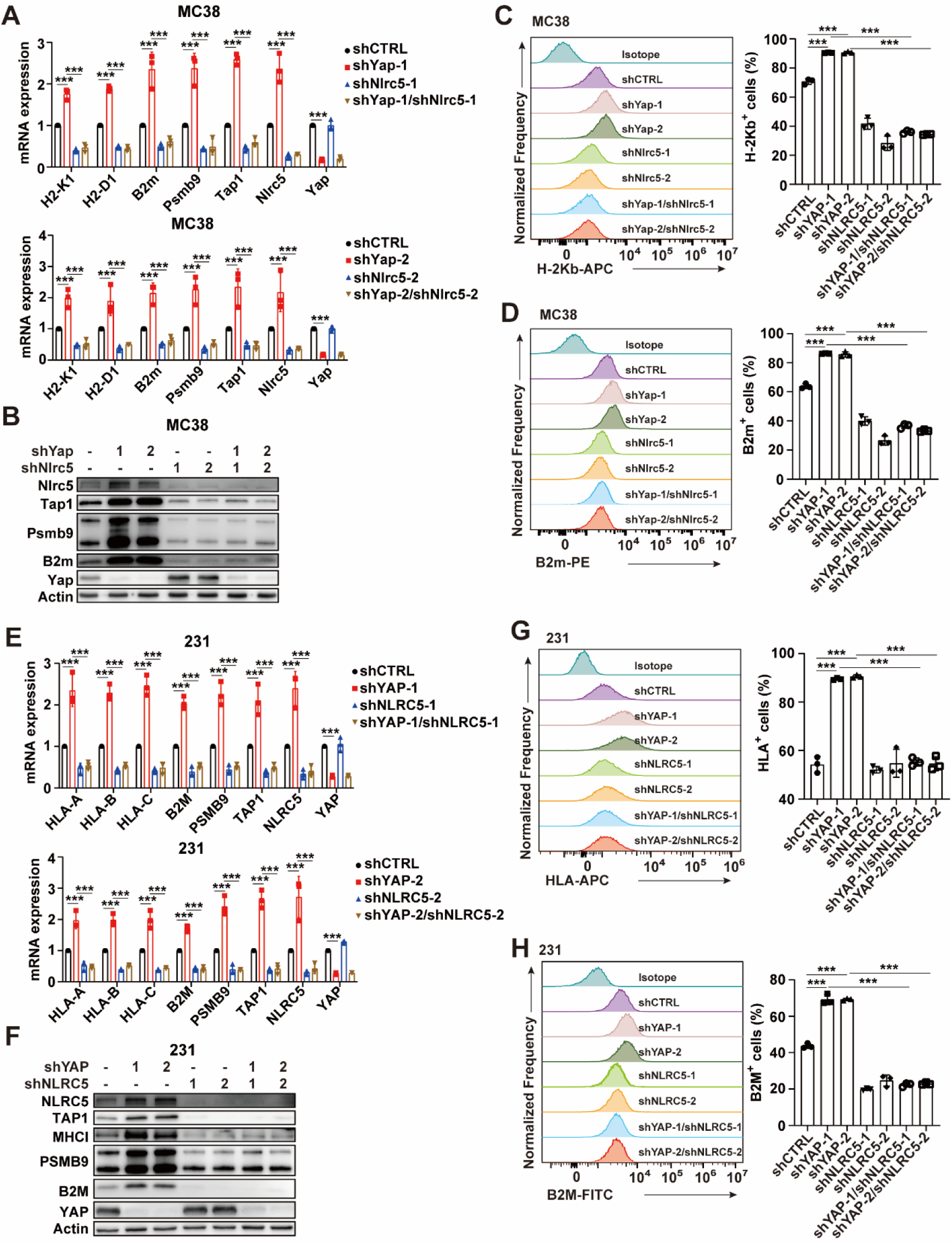
YAP KD induces NLRC5-mediated MHC-I APM gene expression. **(A)** qPCR analysis of the mRNA expression of MHC-I APM genes and Nlrc5 gene in MC38 cells infected with two independent shYap and/or shNlrc5 lentivirus (n = 3). **(B)** Immunoblot (IB) analysis of MC38 cells infected with two independent shYap and/or shNlrc5 lentivirus. **(C)** Flow cytometry detection of cell-surface level of H-2Kb in MC38 cells infected with two independent shYap and/or shNlrc5 lentivirus. The percentage of H-2Kb positive cells is shown (n = 3). **(D)** Flow cytometry detection of cell-surface level of B2m in MC38 cells infected with two independent shYap and/or shNlrc5 lentivirus. The percentage of B2m positive cells is shown (n = 3). **(E)** qPCR analysis of the mRNA expression of MHC-I APM genes and NLRC5 gene in MDA-MB-231 cells infected with two independent shYAP and/or shNLRC5 lentivirus (n = 3). **(F)** IB analysis of MDA-MB-231 cells infected with shYAP and/or shNLRC5 lentivirus. **(G)** Flow cytometry detection of cell-surface level of HLA-A/B/C in MDA-MB-231 cells infected with two independent shYAP and/or shNLRC5 lentivirus. The percentage of HLA-A/B/C positive cells is shown (n = 3). **(H)** Flow cytometry detection of cell-surface level of B2M in MDA-MB-231 cells infected with two independent shYAP and/or shNLRC5 lentivirus. The percentage of B2M positive cells is shown (n = 3). Data are presented as means ± SD. *P* values were calculated by one-way ANOVA in A, B, E and F. *P* values in C, D, G and H were determined by unpaired two-sided *t*-test. \*\*\**P* < 0.001.

**Fig. S7.**
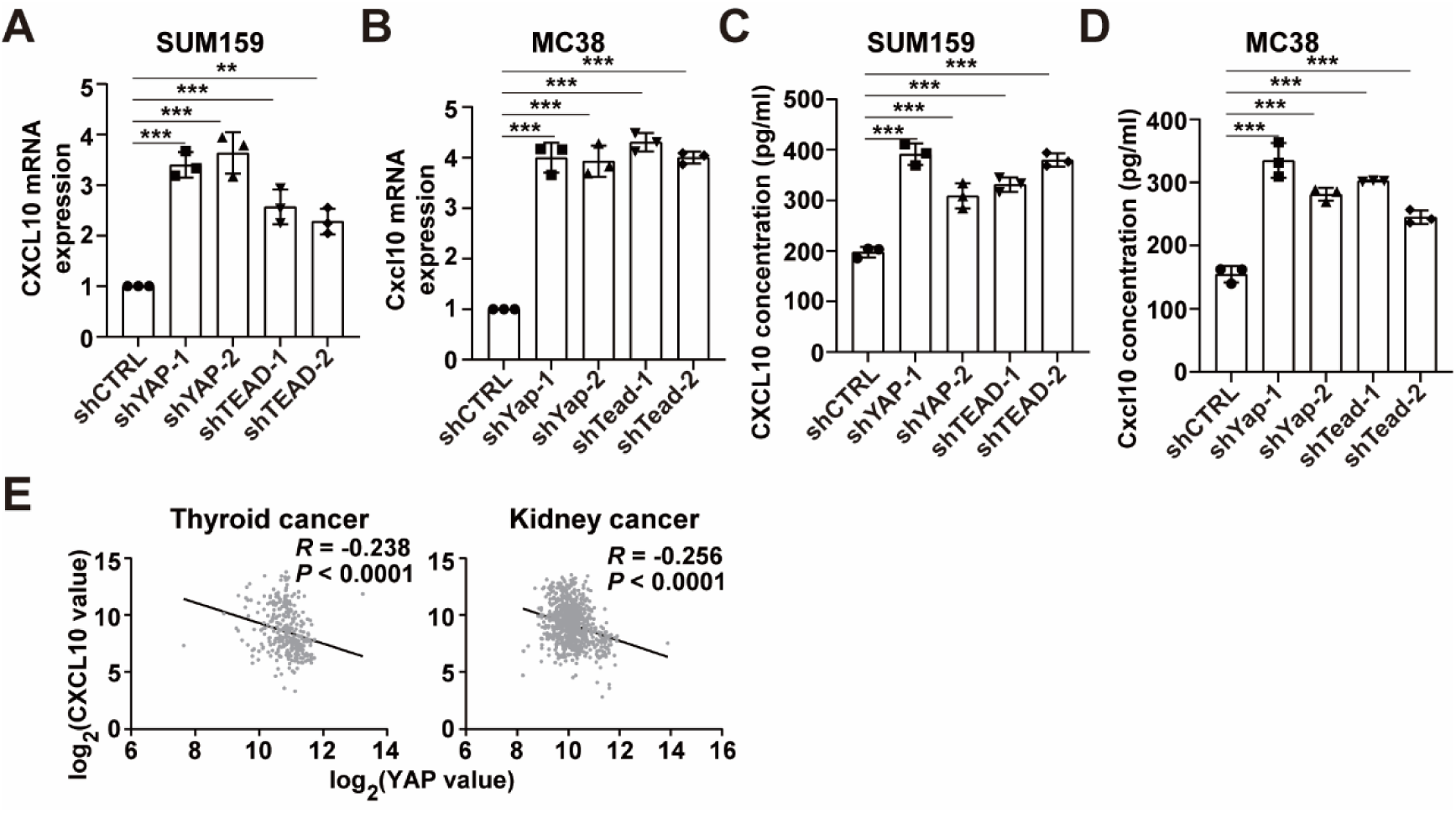
YAP or TEAD KD increases CXCL10 expression. **(A)** qPCR analysis of the mRNA expression of CXCL10 in SUM159 cells with YAP or TEAD1/3/4 KD (n = 3). **(B)** qPCR analysis of the mRNA expression of Cxcl10 in MC38 cells with Yap or Tead2/3/4 KD (n = 3). **(C)** ELISA analysis of CXCL10 protein expression in SUM159 cells with YAP or TEAD1/3/4 KD. **(D)** ELISA analysis of Cxcl10 protein expression in MC38 cells with Yap or Tead2/3/4 KD. **(E)** Negative correlation of YAP mRNA levels with that of CXCL10 in human thyroid cancer (320 samples) and kidney cancer (810 samples). Data from the Gene Expression across Normal and Tumor tissue (GENT) database. *R* represents the Pearson correlation coefficient. Data are presented as means ± SD. *P* values in A, B, C and D were determined by unpaired two-sided *t*-test. *P* value in E was calculated by *F*-test. \**P* < 0.05, \*\**P* < 0.01, \*\*\**P* < 0.001.

**Fig. S8.**
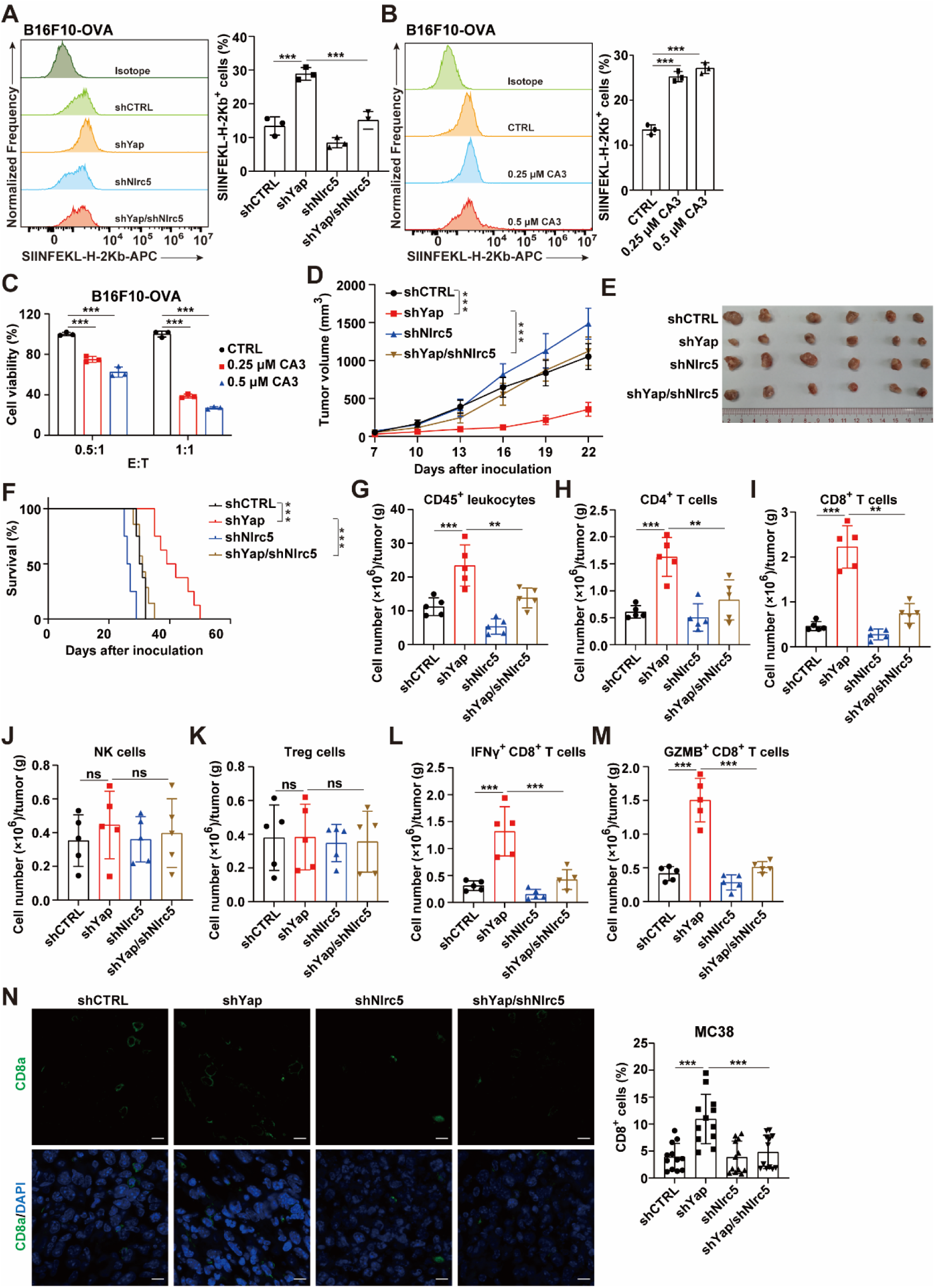
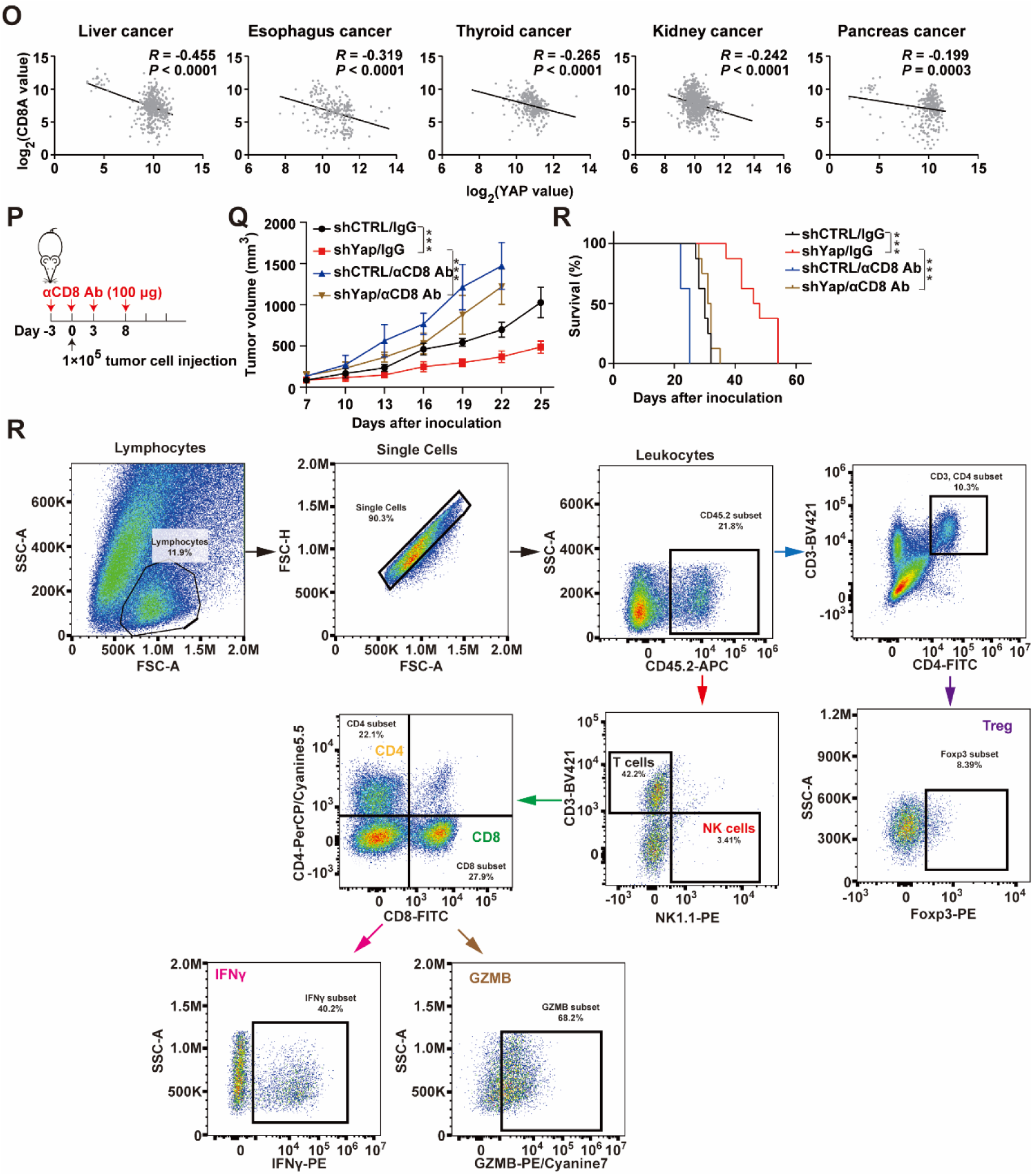
YAP KD promotes intratumoral CTL infiltration via NLRC5. **(A)** Flow cytometry detection of presentation of OVA antigen (SIINFEKL) by MHC I in B16F10-OVA cells with Yap and/or Nlrc5 depletion. The percentage of H-2Kb–SIINFEKL positive cells is shown (n = 3). **(B)** Flow cytometry detection of presentation of OVA antigen (SIINFEKL) by MHC I in B16F10-OVA cells treated with the indicated CA3. The percentage of H-2Kb–SIINFEKL positive cells is shown (n = 3). **(C)** B16F10-OVA cells were treated with the indicated CA3 for 24 h followed by coculturing with OT-1 CD8^+^ T cells for 48 h and the B16F10-OVA cell viability was measured (n = 5). Effector to target (E : T) ratios are shown. **(D to F)** About 1 × 10^5^ mouse MC38 tumor cells with Yap and/or Nlrc5 depletion were subcutaneously (s.c.) inoculated into C57BL/6 mice (n = 8). Tumor size **(D)**, tumor images **(E)** and overall survival **(F)** were monitored. **(G to K)** Flow Cytometric analysis of the indicated immune effector cells in MC38 tumors (n = 5). **(L and M)** Flow Cytometric analysis of IFNγ^+^ and GZMB^+^ CD8^+^ T cells (n = 5). **(N)** Immunofluorescence staining (left) and quantitative estimates (right) of CD8a^+^ cells in MC38 tumours (n = 3). Four fluorescent fields for each of the three samples were counted. Scale bar, 10 μm. **(O)** Negative correlation of YAP mRNA levels with that of CD8A in human liver cancer (517 samples), esophagus cancer (236 samples), thyroid cancer (320 samples), kidney cancer (810 samples) and pancreas cancer (324 samples). Data from the Gene Expression across Normal and Tumor tissue (GENT) database. *R* represents the Pearson correlation coefficient. **(P)** Injection schedule for antibody-mediated depletion of CD8^+^ T cells in C57BL/6 mice. **(Q to R)** About 1 × 10^5^ mouse MC38 tumor cells with or without Yap depletion were subcutaneously (s.c.) inoculated into C57BL/6 mice (n = 8). Tumor size **(Q)** and overall survival **(R)** were monitored. **(S)** Representative flow-cytometry gating strategy for quantifying the numbers of various immune effector cell subsets in murine tumors. Data are presented as means ± SD. *P* values in A, B, C, G, H, I, J, K, L, M and N were determined by one-way ANOVA. *P* values were calculated by two-way ANOVA in D and Q. *P* values in F and R were determined by long-rank test. *P* value in O was determined by *F*-test. \**P* < 0.05, \*\**P* < 0.01, \*\*\**P* < 0.001. ns, not significant.

**Table S1.**
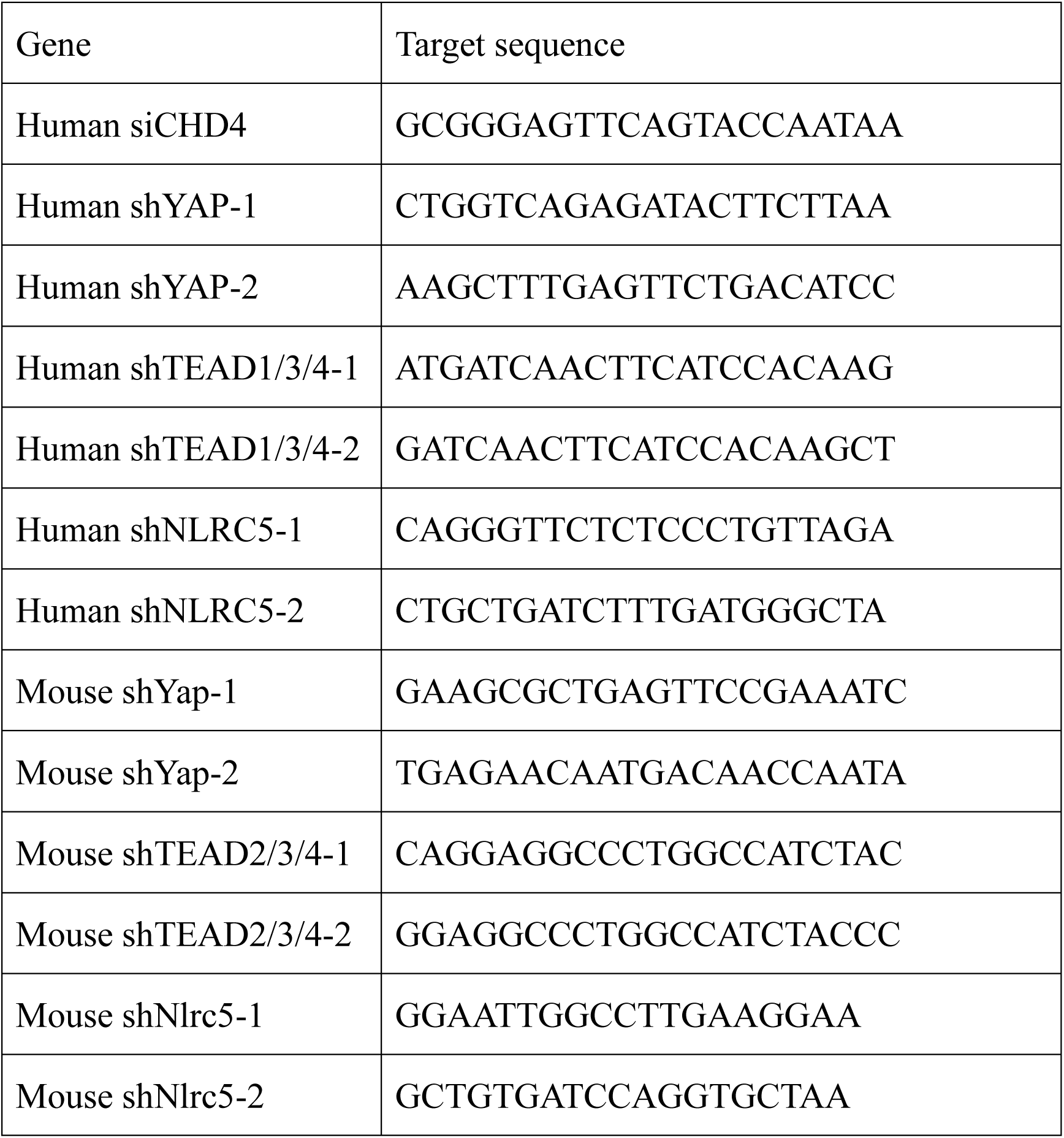
List of siRNA/shRNA in this study.

**Table S2.**
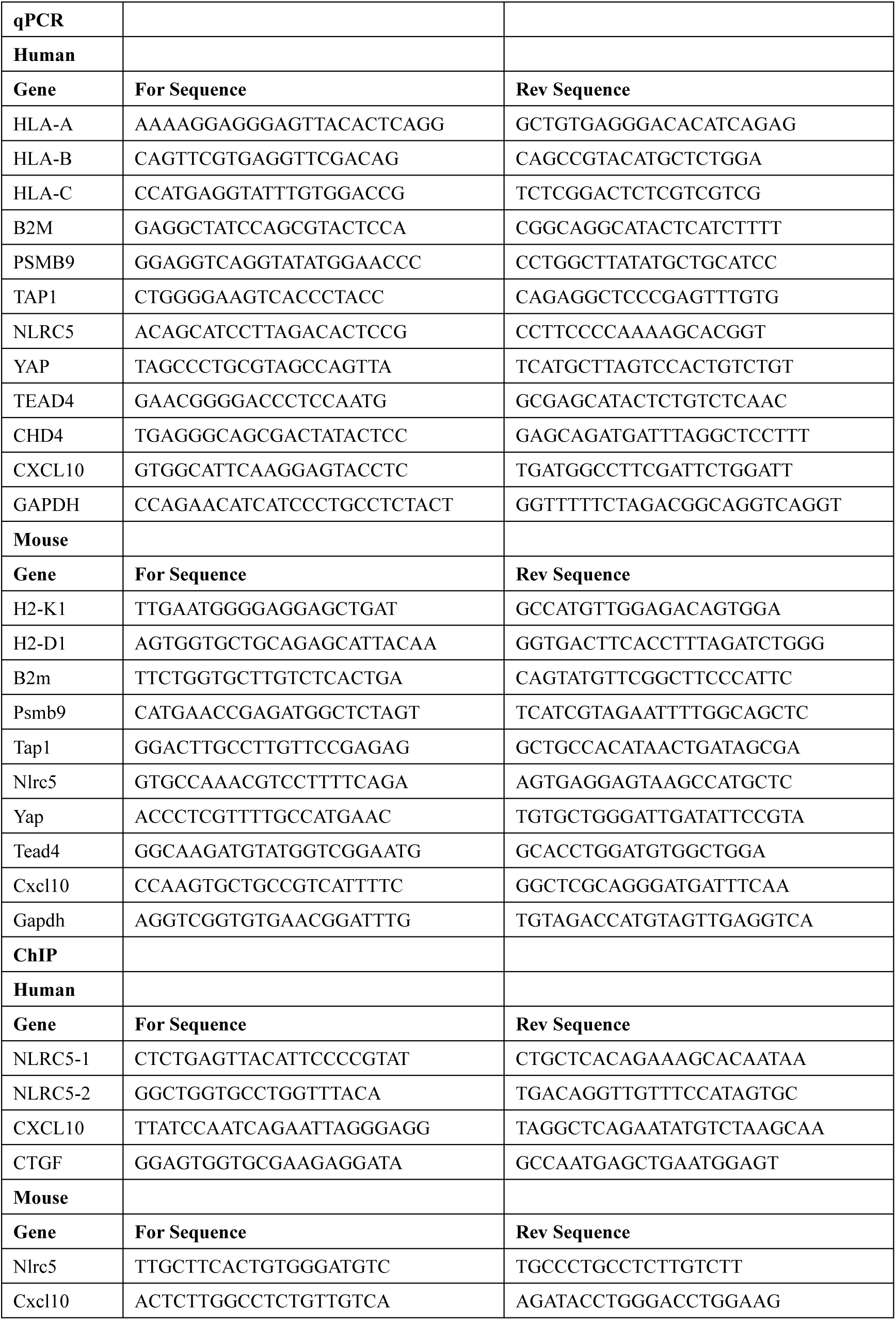
List of primer sequences used in this study.

